# Convergent evolutionary loss of chemosensory and blood-feeding pathways in non-blood-feeding mosquitoes

**DOI:** 10.64898/2026.09.23.753750

**Authors:** Leah Houri-Zeevi, Philipp Brand, Irene Arnoldi, Alexandra E. DeFoe, Jennifer R. Balacco, Nadav Shai, Alex Makunin, Umberto Palatini, Takako Toma, Takao Okazawa, Christopher M. Stone, Anita Schiller, Olivier Fedrigo, Erich D. Jarvis, Mara K. N. Lawniczak, Ichiro Miyagi, Paolo Gabrieli, Leslie B. Vosshall

**Author notes:** Current addresses: Radboud University Medical Center, Nijmegen 6525 The Netherlands (I.A.); Department of Biology and Biotechnology, University of Pavia, Pavia 27100 Italy (U.P.); Colossal Biosciences Inc., Dallas, TX 75226 United States of America (O.F.).

## Abstract

Complex traits that span multiple tissues and systems often integrate large numbers of genes across development, physiology, and behavior, making it challenging to identify their essential components. Blood feeding in mosquitoes is one such trait. It is ancestral to the mosquito family, maintained in most species for ∼200 million years, and was independently lost in three lineages. These convergent losses offer a natural experiment to discover the genetic, physiological, and neural features required for blood feeding. We assembled high-quality, chromosome-level genomes for seven mosquito species, along with whole-brain tomographic reconstructions. Our study spanned the three known non-blood-feeding lineages (*Toxorhynchites rutilus*, *Topomyia yanbarensis*, and *Malaya genurostris*), blood-feeding relatives, and the variable blood feeder *Wyeomyia smithii*. Comparing orthologous gene clades, we detected convergent gene loss specific to the three lineages that had lost blood feeding. The losses include the salivary platelet-aggregation inhibitor *Aegyptin*, blood-activated serine proteases such as Chymotrypsin-1 and 2, and a carboxylesterase expressed in the female fat body and brain glia. The loss of blood feeding also extended to chemosensation. Non-blood feeders lack two odorant-binding protein clades, two ionotropic receptor clades associated with blood-component taste detection, and odorant receptor clades expressed in a discrete, strongly female-biased population of antennal neurons. Female-biased head gene expression was reduced in non-blood feeders. Finally, examination of whole-brain tomographic reconstructions across the species revealed smaller antennal lobes in non-blood-feeding females, consistent with reduced olfactory input. Together, these findings identify a compact set of genes, expression patterns, and brain regions associated with blood feeding, offering an evolutionary entry point for functional dissection of how this complex and dangerous trait is built and dismantled.

## INTRODUCTION

Complex traits such as flight^1^, vision^2^, echolocation^3,4^, vocal learning^5^, and blood feeding^6^ are not the output of a single gene. Instead, they emerge from an integrated system, distributed across many genes, cell types, and organs that must act in concert. This distributed architecture makes the genetic basis of such traits challenging to dissect because a trait spread across dozens of loci and several tissues typically cannot be removed in a single experimental step. Classic forward-genetic screens that disrupt one gene at a time rarely yield phenotypes that offer a comprehensive view of how complex traits are built.

Evolution, which can act on multiple genes simultaneously, offers a powerful alternative approach to understanding complex multi-tissue traits. When the same trait arises independently in separate lineages, this repeated evolution is termed convergent evolution. This repetition functions as a natural experiment, helping to separate new features that are essential for the trait and favored by selection from those that are instead associated with lineage history. Known cases of convergent evolution are widespread. C4 photosynthesis has evolved independently dozens of times among flowering plants^7^. Antifreeze glycoproteins arose separately in Antarctic notothenioid fish lineages and northern cods^8^. Echolocation in bats and toothed whales has converged on the same amino acid substitutions in the inner ear motor protein Prestin^9^. Vocal learning, the rare capacity to imitate sounds, has arisen independently in three distantly related bird lineages, songbirds, parrots, and hummingbirds^5^. It has also emerged in several mammalian lineages, including humans, bats, cetaceans such as dolphins, pinnipeds, and elephants, and it does so through convergence at two levels. One is the independent appearance of dedicated forebrain circuits, a feature that is well documented in songbirds and humans. The other is parallel molecular specializations, including shared regulation of genes such as FOXP2 in the brains of song-learning birds and humans, as well as convergent evolution of other vocal-learning-associated genes and regulatory elements across mammalian vocal learners such as bats and cetaceans^5,10,11^.

The same logic applies when a trait disappears rather than arises. When a trait is abandoned, the genes that built it are freed from the purifying selection that maintained them and they can decay. This way, convergent gene loss across independent lineages informs the mechanism that underlies the trait. This approach has repeatedly connected genotype to phenotype. For example, two geographically distinct cave populations of the Mexican cavefish *Astyanax mexicanus* independently evolved albinism through different loss-of-function deletions in the same pigmentation gene, *Oca2*^12^. The widespread inability to synthesize vitamin C traces to convergent pseudogenization of the same terminal biosynthetic enzyme *GULO* in lineages as disparate as primates, guinea pigs, bats, and passerine birds^13^. Convergent gene loss also informs evolved sensory preferences. Obligate carnivores such as the domestic cat have lost the sweet taste receptor gene *Tas1r2*^14^ and obligate sugar-feeding hummingbirds have repurposed the umami receptor *T1R1-T1R3* to detect sugar^15^. Finally, fully aquatic mammals have shed large fractions of their odorant and taste receptor repertoires^16^. In each case, convergent loss identified the essential genes of a trait.

We were interested in whether the same logic can be extended to the deeply integrated, multi-tissue trait of mosquito blood feeding and whether convergent loss can resolve involvement of important genes in blood feeding. The blood-feeding trait spans sensation, physiology, and reproduction and engages nearly the whole animal. Blood feeding by biting arthropods is a highly specialized trait that demands a coordinated suite of adaptations to locate a host, puncture the skin, avoid host blood clotting and immune defenses, and digest a large, iron-rich protein meal^6^. This behavior has evolved independently across arthropods^17^ and is a rare trait among insects, confined to just a few thousand of the roughly one million described insect species^6^. Yet this small number of blood-feeding arthropod species carries an outsized medical burden because they act as vectors, transmitting pathogens directly into a host as they feed. Collectively they cause many of the world’s most devastating infectious diseases. Mosquitoes transmit malaria, dengue, yellow fever, Zika, chikungunya, lymphatic filariasis, and West Nile; sand flies transmit leishmaniasis; tsetse flies transmit African sleeping sickness; kissing bugs transmit Chagas disease; black flies transmit onchocerciasis (river blindness); fleas and lice transmit plague and typhus; and ticks transmit Lyme disease and babesiosis^18^. Vector-borne diseases account for more than 17% of all infectious diseases and cause ∼700,000 human deaths each year, the vast majority from malaria^18^. Blood feeding is thus a complex trait that is evolutionarily repeated across the tree of life and yet is taxonomically restricted.

Female mosquitoes have fed on blood for approximately 200 million years, and blood feeding is the ancestral state of the mosquito family (Culicidae)^19^. In almost all extant mosquito species, the female must take a blood meal to complete egg development. Seeking a human host recruits a dedicated sensory repertoire that integrates perception of odor, carbon dioxide, heat, and visual cues^20^. Blood acquisition depends on a pharmacologically complex saliva that keeps host blood flowing and blunts immune defenses at the bite site^21^. A single blood meal can exceed the female’s own body weight and triggers a rapid, tightly orchestrated physiological cascade. The meal is concentrated in the midgut where it is proteolytically digested. The free amino acids are used by the fat body to synthesize yolk proteins, which are trafficked to the ovary to mature a batch of ∼100 eggs^22^, and for energetic reserves^23^. A single female can undergo multiple blood-feeding and egg-laying cycles in her lifetime^24^. How such a deeply integrated trait can be assembled or dismantled remains largely unknown.

We reasoned that the rare mosquito lineages that independently abandoned blood feeding offer a natural experiment for defining the core components of this trait. We assembled chromosome-level genomes for seven mosquito species, spanning all three lineages that have independently abandoned blood feeding and their blood-feeding relatives, and asked which genes, cell types, and brain regions were affected by the loss of the trait. We find that non-blood feeders converge on the loss of a compact, shared set of genes involved in saliva, digestion, and reproduction, together with parallel losses in the chemosensory system and brain regions that support host detection. A blood-feeding species nested phylogenetically among the non-blood feeders retains this gene set and neural architecture, indicating that these losses track the trait itself rather than shared ancestry. Together, these results begin to resolve the complex, integrated trait of blood feeding into its constituent molecular and neural parts.

## RESULTS

### Selecting and sourcing mosquito species

We first considered the optimal mosquito species to dissect the genetic and neural basis of blood-feeding loss (Fig. 1a) and concentrated our selection on the Culicinae subfamily of the mosquito tree spanning ∼130 million years of evolution^19^. Among the blood feeders, we examined the yellow fever mosquito *Aedes aegypti*, the invasive and rapidly expanding Asian tiger mosquito *Aedes albopictus*, the frog-biting mosquito *Uranotaenia lowii*^25^, and the iridescent paddle mosquito *Sabethes cyaneus* (Fig. 1b). *Sabethes cyaneus* provides an important phylogenetic control to our study because it is a blood feeder that belongs to the *Sabethini* tribe and is nested phylogenetically among non-blood-feeding genera. Any feature that tracks blood feeding rather than shared ancestry should align *Sabethes* with the blood feeders despite its position in the tree, thereby dissociating the trait from phylogeny.

**Figure 1.**
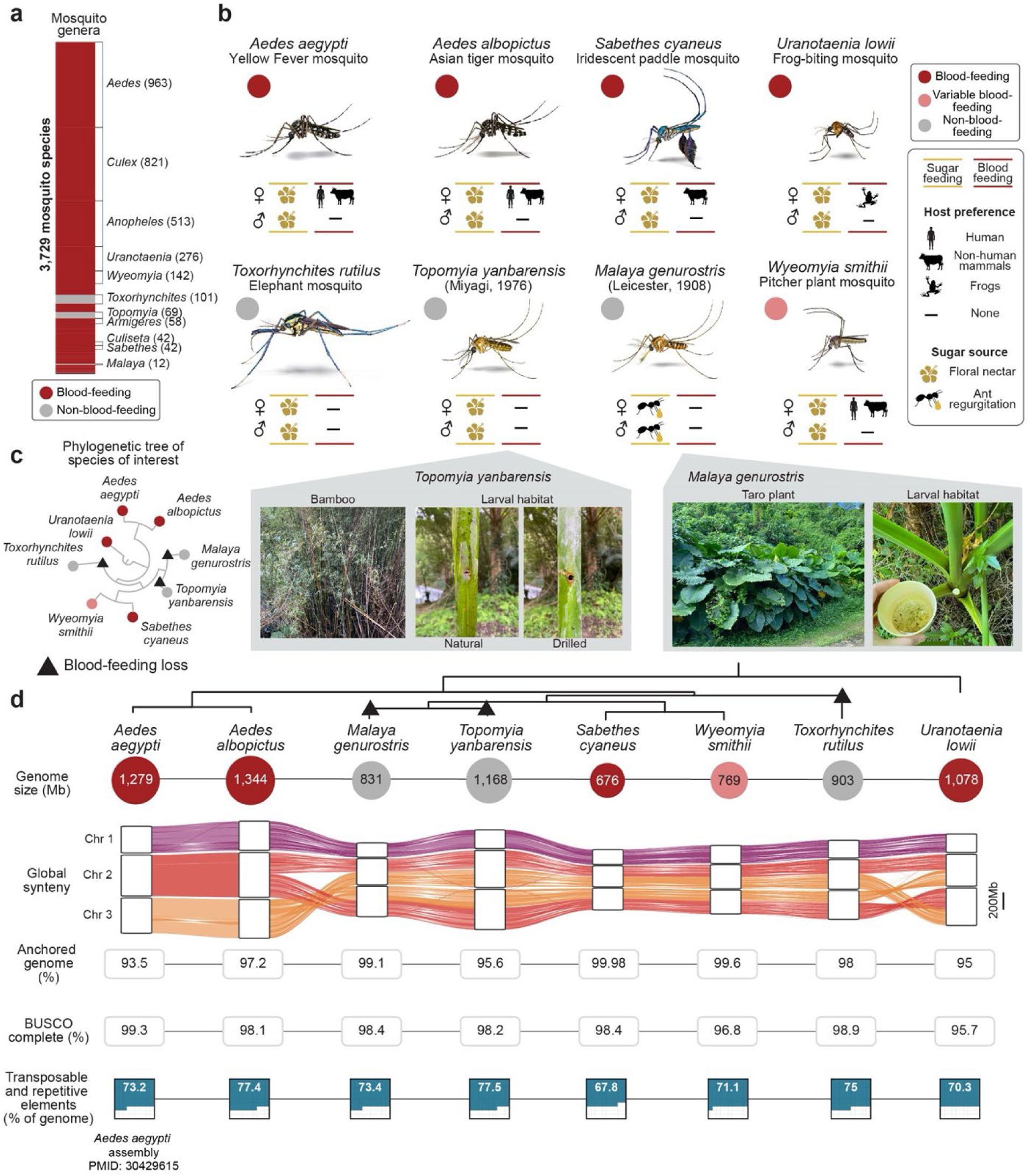
| Species and genomes for the study of blood feeding evolution in mosquitoes **a,** Distribution of blood-feeding (red) and non-blood-feeding (gray) genera across the 3,729 known mosquito species in 42 genera (species counts in parentheses). **b,** Species selected for the study, spanning blood-feeding (dark red), variable blood-feeding (pink), and all known non-blood-feeding (gray) genera. For each species, sex-specific sugar-feeding and blood-feeding behaviors are indicated, together with host preference. A dash denotes absence of the behavior. Bottom: photographs showing natural larval habitats where we collected the two non-blood-feeding species, *Topomyia yanbarensis* and *Malaya genurostris* (bamboo nodes and taro plant leaf axils, respectively). **c,** Phylogenetic relationship among the eight mosquito species, with inferred losses of blood-feeding indicated by black triangles. **d,** Genome assembly statistics for the eight mosquito species, ordered by phylogenetic relationship (blood-feeding losses marked by black triangles). From top to bottom: haploid genome size (Mb); global synteny across the three mosquito chromosomes, with ribbons connecting orthologous genomic blocks between species (scale bar, 200Mb); percentage of the genome anchored to chromosomes; BUSCO completeness scores (%); and transposable and repetitive element content (% of genome). Genome size and phenotype circles are colored as in **b**.

Although some mosquito species can mature a first batch of eggs without a blood meal, they still require blood for subsequent reproductive cycles^24^. To examine true gene loss, we focused our study on the strictly non-blood-feeding mosquitoes, namely, the *Toxorhynchites*, *Topomyia,* and *Malaya* genera. We obtained the elephant mosquito *Toxorhynchites rutilus septentrionalis* (named for its trunk-like proboscis) from an established colony and we field-collected *Topomyia yanbarensis*^26,27^ and *Malaya genurostris* from the island of Okinawa, Japan. Critically, these three lineages, *Toxorhynchites*, *Topomyia*, and *Malaya*, constitute the entire known repertoire of mosquitoes that have lost blood feeding, allowing us to survey the loss of the trait across every lineage in which it has occurred.

Adult mosquitoes of both sexes feed on nectar and other plant sugars for their day-to-day energy needs. In contrast, blood feeding is a female-specific behavior superimposed on this shared sugar diet. Current models suggest that non-blood-feeding lineages acquire the necessary protein reserves for future egg production during development^28^. The aquatic larval stage of *Toxorhynchites* and *Topomyia* species exhibits predatory and sometimes cannibalistic behavior toward mosquito larvae of its own and other insect species. Although *Topomyia* and *Malaya* are closely related, current taxonomic evidence indicates that their losses of blood feeding most likely represent two independent losses (Fig. 1c). The two genera do not form a monophyletic group, and blood-feeding lineages branch between them^19^. Our analysis is agnostic as to whether they reflect one shared or two separate losses of blood feeding. A final species, *Wyeomyia smithii*, occupies a uniquely intermediate position. The species is found in North America and its blood-feeding lifestyle varies by latitude. Some populations blood feed, some do not, and others show mixed phenotypes^29^. Our colonized samples originate from a northern, non-blood-feeding population collected in Ithaca, New York, USA.

### Generation of high-quality chromosome-level genome assemblies

To investigate the genetic basis of blood feeding and its loss, we sequenced and assembled chromosome-level, high-quality genomes for seven of the eight species. Assemblies were generated based on pipelines developed by members of the Vertebrate Genomes Project (VGP), Earth BioGenome Project^30,31^, and Sanger Tree of Life^32^, adjusted for sample preparation of a single insect^33^. All assemblies met the minimum metrics with N50 contig > 1Mb, and QV base accuracy > 40, with three chromosomes for each species. This included resolving to chromosome scale the highly heterozygous genome of the Asian tiger mosquito *Aedes albopictus*, allowing us to build on the knowledge of the previous reference genome^34^. For the eighth species, *Aedes aegypti*, we utilized our published assembly^35^. Assembly sizes ranged from ∼676 Mbp (*Sabethes cyaneus*) to 1,344 Mbp (*Aedes albopictus*). We note that the true genome size of *Aedes albopictus* is estimated at ∼1.25 Gb based on flow cytometry^34^ and the larger value obtained here likely reflects residual haplotypic duplication. The chromosome-level assemblies allowed us to examine global synteny and identify chromosomal arm swaps (Fig. 1d). In each species, 95–99.98% of the assembly was anchored to the main chromosomes, and Benchmarking Universal Single-Copy Orthologs (BUSCO) gene completeness scores ranged from 95.7% to 98.9% (Fig. 1d, Extended Data Fig. 1).

Transposable and repetitive elements accounted for 67.8–77.5% of the genomes (Fig. 1d, Extended Data^36^ – hosted at Zenodo). Endogenous viral elements were integrated at consistent levels across species, with no clear difference between blood feeders and non-blood feeders (Extended Data Fig. 2, Extended Data^36^). Alongside the eight mosquito species described so far we included three more species, *Culex pipiens*, *Culex quinquefasciatus*, and *Armigeres subalbatus*, for which chromosome-level assemblies were already available^37–39^.

### Convergent gene loss in non-blood feeders

What is the core genetic program for blood feeding? To address this, we grouped the protein-coding genes of all species into orthologous clades. Our ortholog-inference approach was sequence-based and phylogenetically aware, performed using OrthoFinder 3^40^. We analyzed the phylogenetic hierarchical orthogroups that correspond to our Culicine group of species and defined blood-feeding-specific gene clades as protein-coding genes that are present in every blood-feeding species and absent from all three lineages that convergently lost blood feeding (Fig. 2a, Extended Data^36^). This criterion trades sensitivity for specificity, and we deliberately favored specificity and applied the filter as stringently as possible, yielding a conservative core set.

**Figure 2.**
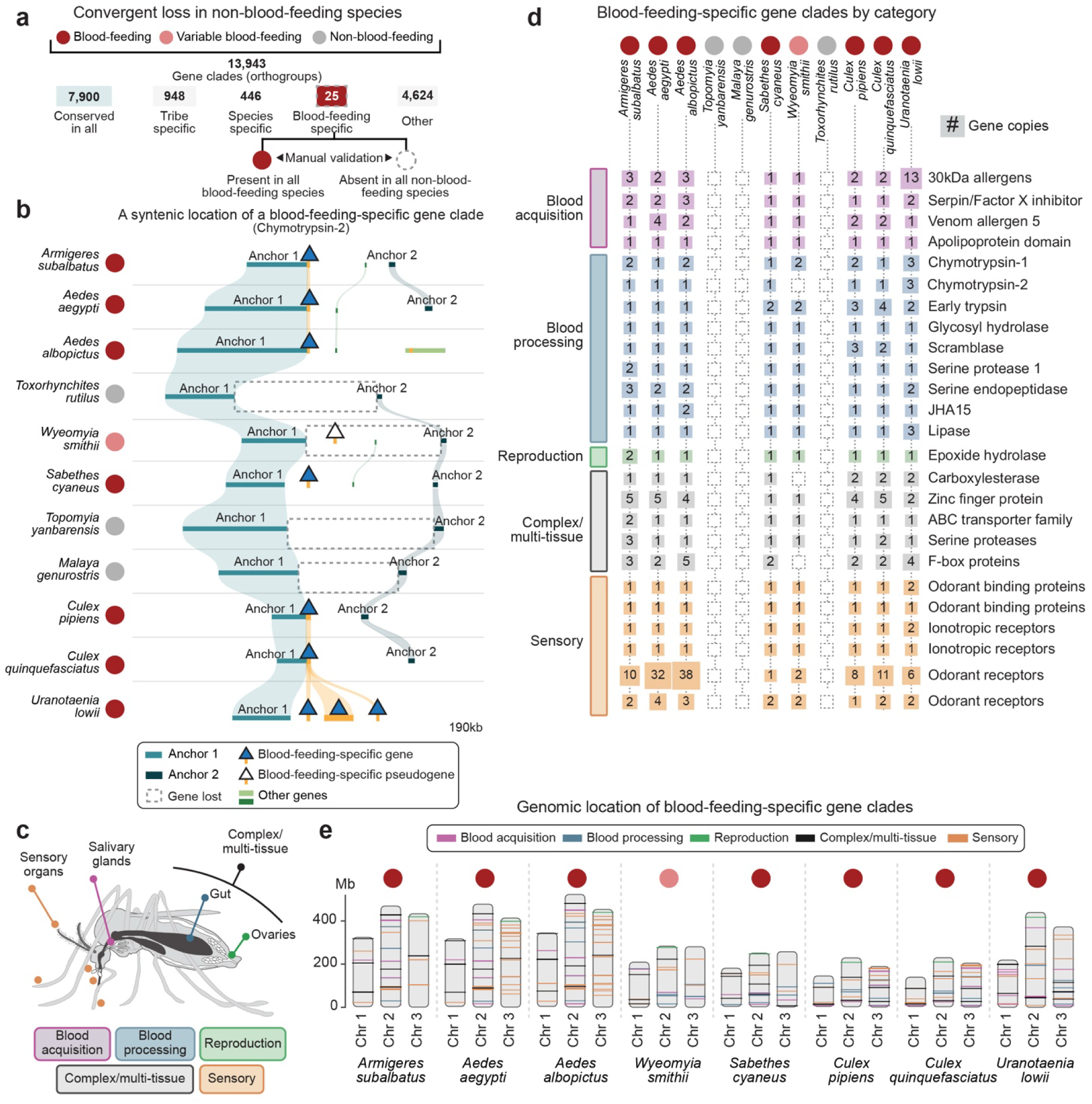
| Convergent loss of blood-feeding-specific gene clades and their functional organization. a, Classification of gene clades (orthogroups) identified across the sampled species. Blood-feeding-specific clades were defined as those present in all blood-feeding species and absent in all non-blood-feeding species and were confirmed by manual validation. **b,** Microsynteny at a representative blood-feeding-specific gene clade (Chymotrypsin-2) across eleven species. Conserved flanking anchor regions (Anchor 1, teal; Anchor 2, dark blue) bracket the locus; the blood-feeding-specific gene (dark blue triangle), pseudogene (open black triangle), gene-loss events (dashed outlines) and other genes (green) are indicated. Region corresponds to 190kb. **c,** Schematic of the functional category and tissue association assigned to blood-feeding-specific genes. **d,** Full representation of each blood-feeding-specific gene clade (rows, grouped by functional category) across species (columns). Numbers and square size indicate the number of gene copies; dashed cells indicate absence. **e,** Chromosomal locations of blood-feeding-specific gene clades across the three chromosomes of blood-feeding and variable blood-feeding species, colored by functional category.

In total we defined 13,943 gene clades. Of these, 7,900 were conserved across all species, 1,394 were tribe– or species-specific, and 26 met the strict criteria for blood-feeding-specific clades. We manually validated these 26 hits and, when applicable, inspected their syntenic positions across genomes to verify true absence rather than annotation or assembly gaps (e.g., Fig. 2b, Extended Data^36^). Manual validation removed two hits and gene-tree inspection added one, yielding a total of 25 blood-feeding-specific gene clades. Combining protein-domain analysis, single-nucleus expression mapping, and a literature survey, we assigned the 25 genes to five broad categories: blood acquisition, blood processing, reproduction, complex/multi-tissue, and sensory function (Fig. 2c, Extended Data Fig. 3). Copy number within these clades varied across species and most remained present in the variable blood feeder *Wyeomyia smithii* (Fig. 2d). In accordance with our filtering criteria, *Sabethes cyaneus*, a blood feeder embedded among non-blood-feeding genera, retained the full set of blood-feeding-specific genes, aligning with the blood feeders despite its phylogenetic position. The blood-feeding-specific genes were distributed across all three chromosomes in the blood-feeding species rather than clustered in a single genomic locus (Fig. 2e).

### Loss of salivary anti-blood-clotting enzymes in non-blood-feeding mosquitoes

We next examined the categories of blood-feeding-specific genes (Fig. 3a), beginning with the blood-acquisition genes (Fig. 3b). Mosquito saliva is a pharmacologically complex secretion produced in the salivary glands, which are organized into distinct lobes with partly specialized secretory functions^41^ (Fig. 3c). Salivary gland transcriptomes of *Aedes aegypti* predict over 100 secreted proteins^42^. As she probes for blood, the female injects this mixture into the host, where the characterized components counteract platelet aggregation, coagulation, and vasoconstriction; dampen inflammation and immune responses; and, in infected animals, facilitate pathogen transmission^43^.

**Figure 3.**
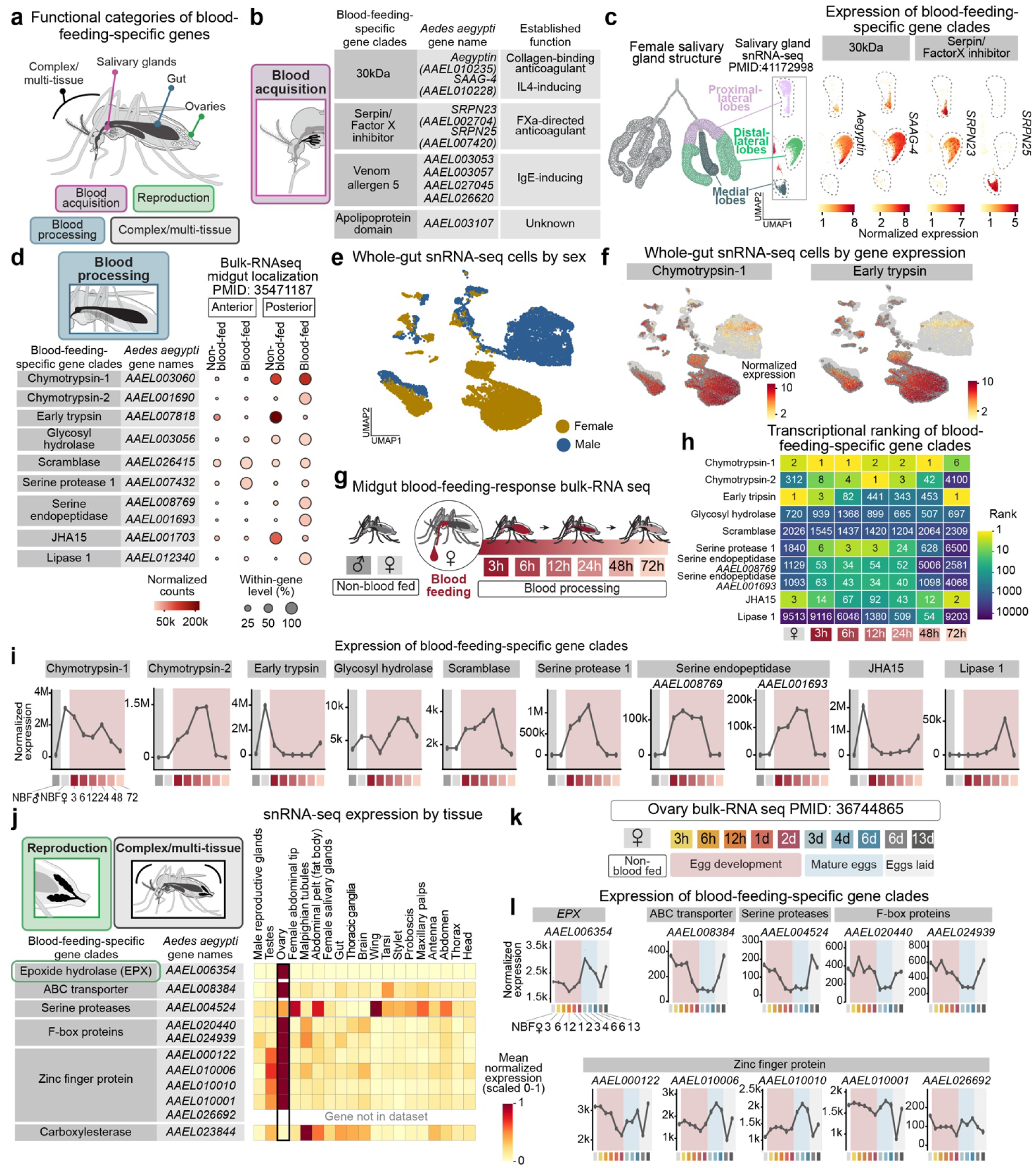
| Blood-feeding-specific genes involved in blood acquisition and blood processing. a, Focus on functional categories and tissue association of blood-feeding-specific genes involved in blood acquisition, blood processing, reproduction, and complex/multi-tissue. **b,** Blood-acquisition gene clades, their *Aedes aegypti* gene identifiers and established functions. **c,** Left, schematic of the female salivary gland showing proximal-lateral, distal-lateral and medial lobes. Right, UMAP projections of salivary-gland single-nucleus RNA-sequencing (snRNA-seq) data colored by normalized expression of representative blood-acquisition genes. **d,** Blood-processing gene clades, their *Aedes aegypti* gene identifiers, and their transcription in anterior and posterior midgut, in sugar-fed female midgut and 48 hours after a blood meal. **e,** UMAP of *Aedes aegypti* whole-gut snRNA-seq cells colored by sex (female, gold; male, blue), representing the sex separation of gut cells in the mosquito. **f,** The gut UMAP colored by normalized expression of Chymotrypsin-1 and Early trypsin. **g,** Design of the midgut blood-feeding-response bulk RNA-seq time course: transcription was examined in midguts of non-blood-fed males and females, and females sampled at 3, 6, 12, 24, 48 and 72 h after a blood meal. **h,** Transcriptional ranking of blood-processing gene clades compared to all other genes across time points (columns), colored by rank (1 = the most highly expressed gene). **i,** Bulk RNA-seq expression trajectories of blood-processing gene clades across the time course (mean); shaded region denotes the male midgut (gray) and the post-blood-meal window (red). **j,** Left, reproduction and complex/multi-tissue gene clades with *Aedes aegypti* identifiers. Right, tissue-level snRNA-seq expression (mean normalized expression scaled 0-1 per gene) across adult tissues. **k,** Design of the ovary bulk RNA-seq time course spanning egg development, mature eggs and egg-laying. **l,** Bulk RNA-seq expression trajectories of reproduction and complex/multi-tissue gene clades across the ovary time course (mean).

We identified four salivary gland-expressed clades. Three have been previously identified as key genes in the female salivary glands and shown to act in blood acquisition and host allergic responses (Fig. 3b)^44–47^. Among them was the gene clade containing *Aegyptin* (*AAEL010235*), an *Aedes aegypti* salivary protein that is known to bind collagen and inhibit platelet aggregation^44^. Recovering these three previously identified blood-feeding-associated proteins purely from convergent loss supports the validity of our approach. The fourth clade encoded an apolipoprotein-domain-containing protein that had not previously been linked to blood feeding. Single-nucleus RNA sequencing (snRNA-seq) of *Aedes aegypti* salivary glands showed that these factors are expressed in defined salivary gland lobes with two blood-feeding serpin paralogues each marking a different lobe (Fig. 3c, Extended Data Fig. 4). The blood-feeding-specific salivary genes mapped almost entirely to the distal-lateral and medial lobes, with only a small, distinct subset of proximal-lateral cells expressing them. These results are consistent with the established functional compartmentalization of the mosquito salivary gland, in which blood-feeding proteins localize to the medial and distal-lateral lobes while the proximal-lateral lobe carries sugar-feeding functions^41,42^.

### Loss of gut blood-processing enzymes in non-blood-feeding mosquitoes

We next turned to genes implicated in blood processing. These clades showed gut-specific expression, and many encoded trypsin– and peptidase-domain proteins. These are serine proteases that cleave dietary proteins into peptides and amino acids and are the principal enzymes that digest the blood meal. Mapping their expression along the gut placed them predominantly in the posterior midgut, which is the compartment responsible for blood digestion in the female (Fig. 3d)^48^. These genes were also markedly female-specific. Chymotrypsin-1 (*AAEL003060*), Early trypsin (*AAEL007818*), and juvenile hormone-regulated serine protease (*JHA15, AAEL001703*) were the top three markers distinguishing female from the non-blood-feeding male in gut enterocytes^49^ [False Discovery Rate (FDR) ∼ 0] and gut cells generally (Fig. 3e-f). To probe their regulation by blood feeding, we carried out bulk RNA sequencing of midguts from sugar-fed males and females and profiled the female midgut transcriptome at several timepoints after a blood meal (Fig. 3g). We discovered several patterns. First, most blood-feeding-specific gut genes were expressed at low levels in males yet were among the most highly expressed transcripts in the female gut before or after blood feeding. The two genes that were expressed in male midgut at high levels, Chymotrypsin-1 and Early trypsin with 100,000 reads each, were expressed at much higher levels in female midguts (3 and 4 million, respectively). Second, each of the blood-feeding-specific gut genes was strongly transcriptionally modulated by blood feeding in the female (Fig. 3h-i). Chymotrypsin-2 and Serine protease 1 were both markedly induced after a blood meal, rising from 13,000 and 2,000 to 1.4 and 1.2 million normalized reads, respectively (Fig. 3i). The Lipase 1 gene showed no meaningful expression (25 normalized reads) in a sugar-fed female (Fig. 3i, Extended Data Fig. 5), and instead was activated post blood feeding, rising to 50,000 normalized reads at 48 hours post blood feeding (2027x fold-change, Fig. 3i).

### Loss of ovary-expressed reproduction and multi-tissue genes in non-blood-feeding mosquitoes

Next, we examined the genes expressed in the ovaries. In mosquitoes, the transition from blood meal to eggs is under endocrine control, with juvenile hormone and the steroid hormone 20-hydroxyecdysone coordinating yolk-protein synthesis and egg maturation across the fat body and ovaries^50^. Consistent with a role in this hormonal axis, we found that one blood-feeding-specific clade encodes an ovary-specific epoxide hydrolase related to the *Drosophila melanogaster* juvenile-hormone epoxide hydrolase, an enzyme that inactivates juvenile hormone (Fig. 3j, Extended Data Fig. 6)^51^.

We also identified five clades of genes that could not be assigned to a single tissue or function, in keeping with the multi-systemic nature of blood feeding, and investigated their cellular expression. We refer to these as complex/multi-tissue. Among these genes are an ABC transporter (*AAEL008384*) that is expressed in multiple blood-related tissues and cell types including the female germline and follicles, a blood-feeding-specific antennal neuronal cluster cell type, and *Ir25a* neurons in the tarsi and proboscis. The expression of the ABC transporter marked neuropeptide-F-expressing gut enteroendocrine cells [log fold-change = 3.9, adjusted p-value (p adj) = 2.4 x 10^−11^] (Extended Data Fig. 6). We identified a blood-feeding-specific CLIP-domain serine protease (*CLIPC5B*, *AAEL004524*) expressed in the female follicle and epithelial cells across tissues. Several CLIP-domain serine proteases were previously shown to mediate mosquito and insect innate immunity^52,53^. Another blood-feeding-specific gene is the *CCEae3A* carboxylesterase (*AAEL023844*) (Fig. 3j), previously implicated in insecticide resistance^54^. This gene shows strong expression in the female fat body and brain glia (Extended Data Fig. 6), two tissues that are key for the blood-feeding cycle and hormonal responses^22,49^.

Because most of these genes are strongly transcribed in the ovaries, we examined their expression through the ovarian response to blood feeding and across the egg-development and egg-laying cycle^55^. We found that the expression of these genes is regulated in a blood-feeding– and oviposition-cycle-dependent manner (Fig. 3k-l). Specifically, both the ABC transporter and *CLIPC5B* were strongly down-regulated one day after blood feeding and as long as mature eggs were present, returning to baseline after egg laying. In contrast, epoxide hydrolase and two zinc finger proteins showed the reverse pattern and were instead upregulated upon egg maturation (Fig. 3l).

### Convergent loss of chemosensory genes in non-blood feeders

The female mosquito depends on a sensitive and integrative sensory system to find and evaluate potential hosts. She combines perception of host odor, exhaled carbon dioxide, and body heat with visual contrast to orient toward a host. Once she has punctured the skin, she can sense adenosine-5′-triphosphate (ATP) and other blood components to decide whether to feed (Fig. 4a)^56–58^. We asked which sensory genes and modalities are specifically tied to blood feeding and identified convergently lost clades of odorant-binding proteins, ionotropic receptors, and odorant receptors. Each was strictly conserved in blood feeders and absent from all non-blood feeders (Fig. 4b).

**Figure 4.**
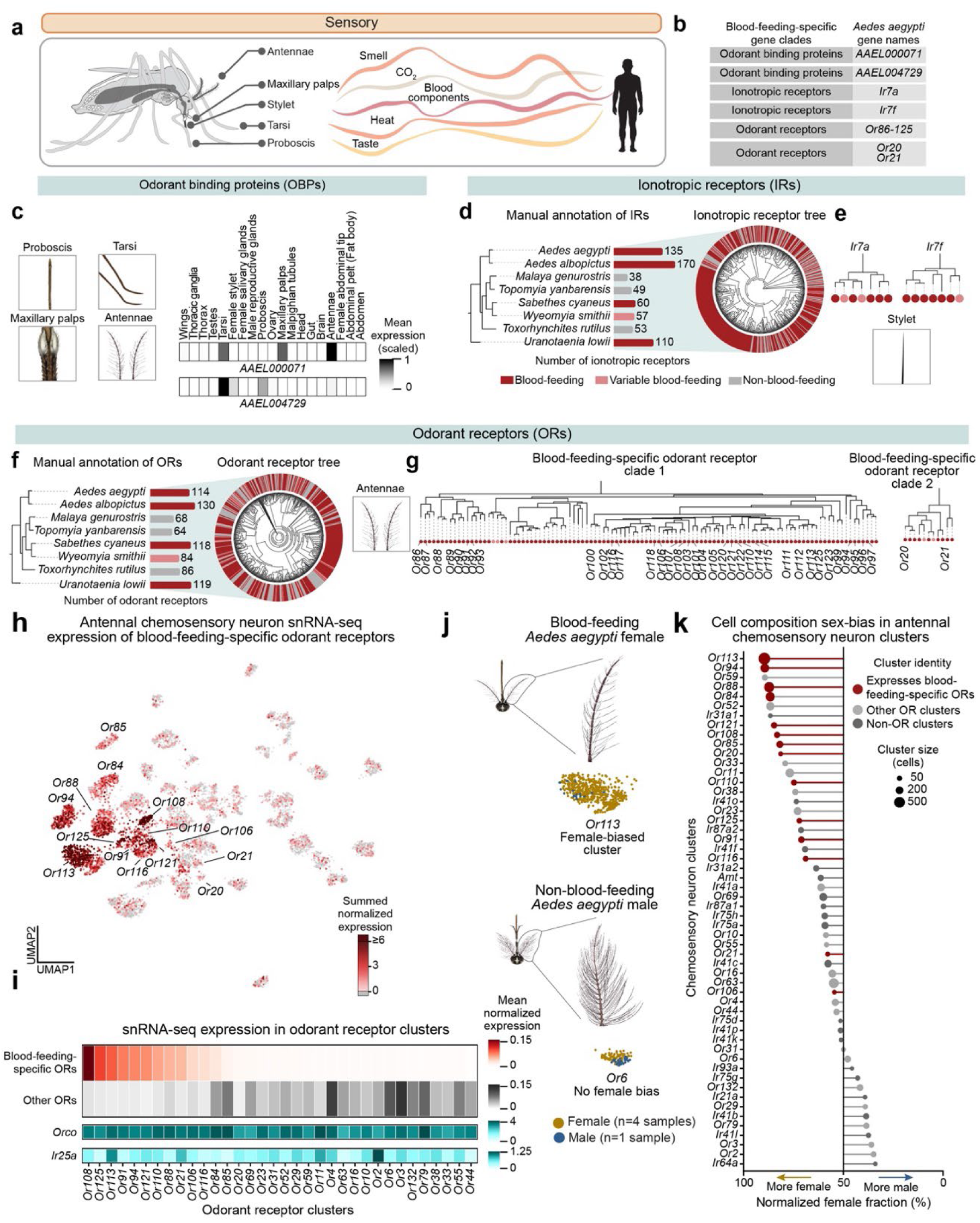
| Sensory blood-feeding-specific gene clades and their cellular expression a, Schematic of the mosquito sensory appendages involved in the detection of host-derived cues. **b,** Sensory blood-feeding-specific gene clades and their *Aedes aegypti* gene identifiers. **c,** Blood-feeding specific odorant-binding proteins (OBPs). Left, representative sensory appendages. Right, tissue expression (mean expression, scaled 0-1) of *AAEL000071* and *AAEL004729* across adult tissues. **d,** Ionotropic receptors (IRs). Left, manually annotated IR counts per species mapped onto species phylogeny (bars colored by phenotype). Right, gene tree of all IRs with the outer ring colored by the species feeding state. **e,** Gene trees for the stylet-expressed blood-feeding-specific IR clades *Ir7a* and *Ir7f*, with tip circles colored by species feeding state. **f,** Odorant receptors (ORs). Left, manually annotated OR counts per species (as in **d**). Right, all ORs gene tree with the species feeding state represented by the outer ring. **g,** Blood-feeding-specific OR clade 1 (*Aedes aegypti Or86-Or125*) and clade 2 (*Aedes aegypti Or20*, *Or21*) branches of the gene tree. **h,** UMAP of antennal chemosensory-neuron snRNA-seq data colored by summed normalized expression of blood-feeding-specific ORs. Blood-feeding specific OR-defined clusters are labeled. **i,** Heatmap of mean normalized expression across OR clusters (columns) for blood-feeding-specific ORs and other ORs, with *Orco* and *Ir25a* shown below. **j,** Representative antennal morphology and OR-defined clusters for blood-feeding *Aedes aegypti* female (*Or113*, female-biased cluster) and non-blood-feeding *Aedes aegypti* male (*Or6*, no-female-bias cluster); female, *n* = 4 samples; male, *n* = 1 sample. **k,** Sex bias in antennal chemosensory-neuron cluster composition, ordered by normalized female fraction (%). Circles are colored by cluster identity (blood-feeding ORs, dark red; other OR clusters, gray; non-OR clusters, dark gray) and sized by cluster size (cells). Clusters expressing blood-feeding-specific ORs (dark red) are those defined by a blood-feeding-specific OR (Or86-Or125, Or20, Or21) or, for Or84 and Or85, clusters defined by a retained OR that show high summed expression of blood-feeding-specific ORs (Fig. 4h,i).

Insect chemosensory neurons are housed in specialized cuticular structures called sensilla that are bathed in aqueous sensillum lymph. Odorant-binding proteins (OBPs) are small, abundant proteins typically secreted from epithelial support cells and thought to partition hydrophobic odorants from the air to the lymph where they can interact with chemosensory receptors^59^. The two blood-feeding-specific OBPs were expressed in key sensory tissues in the mosquito. *AAEL000071* was expressed in tarsi, maxillary palps, and antennae, and *AAEL004729* was expressed in proboscis and tarsi (Fig. 4c). *AAEL000071* showed the expected epithelial-like cell expression (*snu*-positive, Extended Data Fig. 7a). Unexpectedly, the other odorant-binding protein, *AAEL004729*, was specifically and significantly expressed in *prospero-NompA* cell clusters and in *Ir25a* and *Gr7* neuronal clusters in both the mosquito proboscis (*prospero-NompA*, log fold-change = 4.9, p adj < 3.9 x 10^−18^; *Ir25a* neurons, log fold-change = 3.1, p adj < 1.4 x 10^−17^; Gr7 log fold-change = 3.2, p adj < 1.8 x 10^−8^) and tarsi (*prospero-NompA*, log fold-change = 8.8, p adj < 2 x 10^−83^; *Ir25a* neurons, log fold-change = 4.8, p adj < 2 x 10^−83^; *Gr7* log fold-change = 1.8, p adj < 1.6 x 10^−6^) (Extended Data Fig. 7b-c). We do not have insights into the function of either OBP or why their cell-type expression differs.

Insect chemoreceptors evolve rapidly, with frequent gene birth, death, and lineage-specific expansion that track ecological shifts in diet and host preference^60^. There are three major families of chemosensory genes that detect sensory stimuli in the mosquito. Ionotropic receptors (IRs) are evolutionarily related to ionotropic glutamate receptors and detect a diversity of stimuli including acids, amines, humidity, and temperature^61–63^. Odorant receptors (ORs) are an insect-specific gene family that, together with the co-receptor Orco, detect volatile odors^64,65^. Gustatory receptors (GRs) are an ancient chemoreceptor family from which ORs evolved, and mediate taste, CO_2_ detection, and contact pheromone sensing^20,66^.

To map the chemoreceptor repertoire accurately across species, we manually annotated GRs (Extended Data Fig. 7d, Extended Data^36^), IRs (Fig. 4d, Extended Data^36^), and ORs (Fig. 4f, Extended Data^36^). We found no blood-feeding-associated difference in GR copy number or convergent loss (Extended Data Fig. 7d, Extended Data^36^). IR counts were substantially higher in *Aedes* and *Uranotaenia* species compared to the other genera, due to a large lineage-specific IR expansion (Fig. 4d, Extended Data Fig. 7e). Because IR number was low in the non-blood feeders but equally low in their close blood-feeding relative, *Sabethes cyaneus*, we categorized these differences as phylogenetic rather than feeding mode.

Beyond these numerical differences, however, we identified two blood-feeding-specific IR clades, *Ir7a* and *Ir7f* (Fig. 4e). These genes were previously shown to be specifically expressed in the female stylet in neurons that are activated in response to stimulation by blood and blood components^56^.

### Blood-feeding-specific odorant receptors define a distinct population of female antennal neurons

We further examined the odorant receptors, a gene family required for the detection of many host-derived volatiles and for discrimination of human from non-human animal odor^67^. Rather than following phylogeny as we found for most IRs, total OR number tracked the blood-feeding trait (Fig. 4f). Each non-blood-feeding lineage, together with the variable blood feeder *Wyeomyia smithii*, encoded a reduced OR repertoire (64-86 total ORs), whereas every blood feeder, including *Sabethes cyaneus,* retained a large one (114-130 total ORs). Asking which clades had been lost, we identified two blood-feeding-specific OR clades, corresponding to *Aedes aegypti Or86-Or125* and *Or20-Or21* (Fig. 4g, Extended Data Fig. 7f). As with the other blood-feeding-specific genes, these were not confined to a single genomic locus (Fig. 2e). Mapping their expression in the *Aedes aegypti* antenna snRNA-seq dataset, we found that these receptors define a discrete subset of chemosensory neurons in the antenna (Fig. 4h), and that the remaining, non-blood-feeding-specific ORs were rarely co-expressed in those clusters (Fig. 4i). Because blood feeding is a female-specific behavior, we asked whether these clusters were sexually dimorphic (Fig. 4j). We found that the cell composition of neuronal clusters expressing blood-feeding-specific ORs was strongly female-biased, accounting for 8 of the 10 most female-biased OR clusters, including the two large and most female-biased clusters overall, *Or113* and *Or94* (Fig. 4k). Together, these results reveal the convergent loss of multiple components of the mosquito sensory system upon the loss of blood feeding and pinpoint a set of blood-feeding-specific odorant receptors expressed in a distinct, strongly female-biased population of neurons.

### Convergent reduction of female pharyngeal muscles and female-biased head gene expression

Across the blood-feeding-specific genes, we repeatedly encountered female-specific or female-biased expression. We therefore set out to examine female bias directly, in both internal head anatomy and in gene expression (Fig. 5a-c). To do so, we used propagation-based phase-contrast synchrotron microtomography (synchrotron micro-CT) to scan and reconstruct the internal head and brain anatomy of each of the eight species studied here and of additional blood-feeding mosquitoes (Fig. 5b, Extended Data Figs. 8, and 10). The non-neural internal head structures revealed a clear enlargement of female pharyngeal muscles compared to the male. These muscles power the female’s pharyngeal pump, which is part of her blood-sucking machinery. This sexually dimorphic muscle size was strongly present in the blood-feeding species, and the pharyngeal muscles of blood-feeding females were significantly larger than those of non-blood feeding females (q-value =0.045) (Fig. 5b, Extended Data Fig. 8a-b).

**Figure 5.**
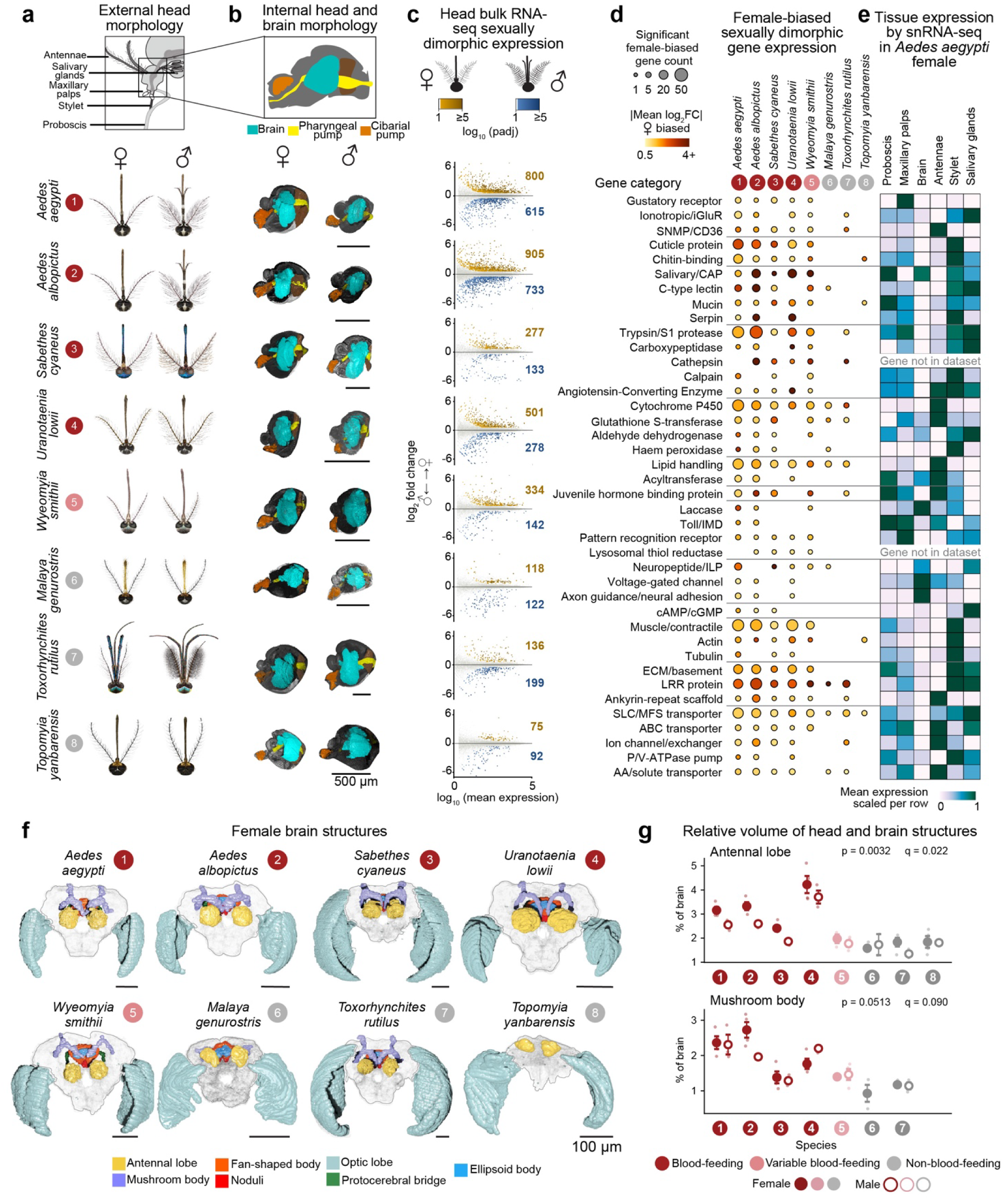
| Sexually dimorphic head morphology, gene expression and neuroanatomy across species. a, Illustrations of the external head morphology of females and males for the indicated species (numbered 1-8; blood-feeding phenotype color as in Fig. 1). **b,** Internal head and brain morphology. Scale bar, 500 µm. **c,** MA plots of head bulk RNA-seq sexually dimorphic expression per species, with significantly female-biased (gold) and male-biased (blue) genes; total counts of female– and male-biased genes are indicated (FDR<0.05, FC>=1.5). **d,** Female-biased sexually dimorphic gene expression by functional gene category (rows) across species (columns; phenotype circles as in Fig. 1). Circle size denotes the number of significant female-biased genes and color the mean absolute log_2_ fold change. **e,** Tissue expression of the same gene categories by snRNA-seq in *Aedes aegypti* female across head tissues (mean expression scaled 0-1 per row). **f,** Segmented female brain structures for each species, colored by neuropil. Scale bar, 100 µm. Only antennal and optic lobes were segmented for *Topomyia yanbarensis* due to sample quality. **g,** Relative volume (% of total brain) of the antennal lobe and mushroom body (numbered as in **f**), shown for females (filled) and males (open). Points show individual measurements; main symbols denote the mean.

To quantify sexually dimorphic gene expression, we sequenced bulk RNA from male and female heads across the mosquito species (Fig. 5c). Transcriptionally, blood-feeding species had more sex-differentially expressed genes overall and more female-biased genes compared to non-blood feeders (Fig. 5c, Extended Data^36^). To identify the pathways underlying this reduced female bias, we grouped the differentially expressed genes by functional category using domain and functional inference (Fig. 5d, Extended Data Figs. 8c, and 9). Female bias in blood feeders spanned several categories, including three *Aedes aegypti* gustatory receptors *Gr1*, *Gr2*, and *Gr3*, which are expressed in the maxillary palps and together constitute the carbon dioxide receptor complex^20^, as well as cuticular proteins, salivary proteins, and muscle-related proteins (Fig. 5d). We further mapped these genes across head cell types to anchor the expression bias to specific structures and discovered female– and blood-feeding-biased transcriptional categories which are strongly present in brain, antenna, and stylet tissues, as well as female salivary glands (Fig. 5e).

### Loss of blood feeding is associated with smaller olfactory processing centers in the brain

Sensory input converges on the brain, and the peripheral differences we observed led us to ask whether brain anatomy differs according to blood-feeding phenotype. We therefore reconstructed whole-brain morphological atlases across blood-feeding and non-blood-feeding species (Fig. 5f, Extended Data Fig. 10, Extended Data^36^) and asked which regions differ between the two feeding states. For *Topomyia*, the available material only allowed confident reconstruction of the antennal and optic lobes, while complete internal atlases were resolved for the remaining species. We found the antennal lobes, the primary olfactory centers of the mosquito where olfactory sensory neurons synapse onto projection neurons within discrete glomeruli, were significantly reduced in non-blood–feeding females (1.9-fold, PGLS, p = 0.003, q = 0.022) (Fig. 5g, Extended Data Fig. 10). The mushroom bodies, a brain region that specializes in olfactory learning and sensory integration and one of the two principal targets of antennal lobe projection neurons^68^, showed a consistent reduction in non-blood-feeders compared to blood-feeders, although it did not reach significance cutoff (1.8-fold, PGLS, p = 0.051, q = 0.090) (Fig. 5g, Extended Data Fig. 10). Antennal lobe size was related to feeding phenotype rather than shared ancestry: *Sabethes cyaneus*, the embedded blood-feeding control, retained blood-feeder-like antennal lobes despite its position among non-blood-feeding genera. *Wyeomyia smithii*, the variable blood feeder, which was held out of the model, had antennal lobes reduced to the size seen in non-blood feeders (Fig. 5g).

## DISCUSSION

Blood feeding is a complex trait, involving sensation, digestion, reproduction, and hormonal regulation. By exploiting the convergent loss of blood feeding in mosquitoes, we defined a set of genes, expression patterns, and brain regions that are associated with blood feeding. Our approach recovered canonical blood-associated factors such as *Aegyptin*, a salivary platelet-aggregation inhibitor, and posterior-midgut digestive proteases. It also revealed previously unsuspected players, including an apolipoprotein-domain salivary protein, an ABC transporter, a CLIP-domain serine protease, a neuronally expressed odorant binding protein, and a specific subset of odorant receptors. It is interesting that the convergent loss of blood feeding did not surface a clear subset of immune-related proteins in the mosquito, perhaps hinting at the rapid evolution of the immune system.

The convergent shrinkage of the antennal lobes mirrors the peripheral loss of odorant receptors, suggesting a reduced sensory input that is accompanied by a coordinated reduction of the central circuits that process it. Whether these volumetric reductions reflect fewer neurons, sparser connectivity, or reduced neuronal arborization remains to be determined. The convergent reduction of these structures in non-blood feeders could suggest that these brain tissues track the sensory demands of finding and feeding on a host.

*Wyeomyia smithii*, whose geographically distributed populations span the transition from blood feeding to non-blood feeding, offers a rare window into the process of blood feeding loss. It retained most blood-feeding-specific genes yet has a contracted odorant receptor repertoire and reduced antennal lobes. If read as a snapshot of the loss of a complex trait in progress, this mosaic suggests a sequence in which the odorant receptor repertoire and its central targets contract before the peripheral effector genes of blood acquisition and digestion are erased. Establishing the true order, whether there is a genetic switch for blood feeding plasticity in these animals, and whether one change drives the next, will require targeted mechanistic studies.

The reach of blood feeding extends beyond genes, neurons and brain regions mapped here. In a companion study^69^ we report that blood feeding is mirrored in the mosquito proteome itself. The amino acid composition across the proteome of blood-feeding mosquitoes converged on lower usage of isoleucine, reflecting mammalian hemoglobin amino acid bias. The restriction of isoleucine in adult hemoglobin in turn may have its origins in the blood-borne parasites that mosquitoes harbor and transmit. Read together, the two studies frame blood feeding as a trait that has left its imprint at every level of the animal, from the composition of its proteins to the regions of its brain.

What allows a lineage to abandon blood feeding in the first place? One likely prerequisite is an alternative route to the protein that eggs would otherwise draw from blood. Indeed, in the variable blood feeder *Wyeomyia smithii*, the female’s ability to reproduce without consuming a blood meal has been linked to larval nutrition and environment^70^. The three lineages studied here further hint at such solutions. *Toxorhynchites* and *Topomyia* may meet this demand through larval predation. Their larvae are avid, sometimes cannibalistic predators of other mosquito and insect larvae, a trait that has made *Toxorhynchites* of interest as a biological control agent against mosquito vector species^71^. The feeding behavior of *Malaya genurostris* is perhaps the most peculiar. While its larval nutrition remains to be fully understood, its adults feed on neither blood nor nectar. Instead, both males and females of this species use a distinctively swollen, apically expanded proboscis to stimulate ants to regurgitate. The mosquitoes then feed on ant-derived honeydew, the sugary liquid ants collect from sap-feeding insects and regurgitate, in a mode of kleptoparasitism^72–74^. The *Sabethini* tribe might offer an additional clue. This tribe includes the non-blood feeders *Topomyia* and *Malaya* and the variable feeder *Wyeomyia smithii*. Interestingly, several Sabethini species have predatory larvae, and *Sabethes cyaneus* larvae are occasional predators of other mosquito and insect larvae^75^ which could suggest a predisposition in this tribe for a reduced dependency on, and eventual loss of, blood feeding. In each of these cases, a shift in feeding strategy could supply reproductive resources without host contact and thereby relax the selection that maintains the blood-feeding trait. Whether such shifts are causes of the loss of blood feeding or consequences of it, and what genetic or ecological changes permit or promote them, remains to be established.

Finally, by tracing blood feeding through the species that abandoned it, this study provides an entry point into the molecular and neural evolution of this complex, medically important trait. The genes, expression patterns, and brain structures we find associated with mosquito blood feeding provide concrete targets for dissecting this behavior at higher resolution. Future work will need to investigate the processes that sustain or dismantle blood feeding and more broadly, how complex traits are assembled and disassembled by evolution.

## LIMITATIONS OF THE STUDY

By design, this study revealed a correlation of genes, expression patterns, and brain regions associated with the loss of blood feeding. Our work does not establish that any of these are causally required for the trait. Future work using CRISPR-based genome editing to disrupt candidate genes in blood-feeding species would be needed to clarify whether aspects of the trait can be dismantled by reverse genetics. Another approach would be to reintroduce candidate genes into the genomes of non-blood-feeding mosquitoes to determine sufficiency for any aspect of the complex blood-feeding trait. Both approaches will need to bear in mind that the redundancy built into mosquito physiology may buffer the loss or gain of any single component^20,67,76,77^.

Our gene clade loss criterion was deliberately stringent, and the 25 blood-feeding-specific gene clades reported here likely represent a conservative subset of the total possible genetic units associated with blood feeding. Additional genes, specific amino acid changes, and changes in gene regulatory sequences are likely to be relevant as well. Although our genomic sampling covered every known non-blood-feeding lineage, it remains a limited selection of mosquito species diversity and new genomes will be needed to test how far these findings generalize. Finally, the synchrotron micro-CT reconstructions capture gross morphology only and the sample size was limited. The anatomical differences we describe will need further resolution at the cellular and circuit level to understand how the brain of the blood-feeder differs from the non-blood feeder.

## ACKNOWLEDGMENTS

We thank members of the Vosshall Lab for discussion and comments on the manuscript; Gloria Gordon and Libby Mejia for strain maintenance; Marta Villa and Laura Soresinetti for help with analysis of the head synchrotron scans; Alan Tracey, Jonathan Wood, Katharina von Wyschetzki, and Shane A. McCarthy for assistance on the *Sabethes cyaneus* genome; Mr. T. Ganaha and Ms. A. Chien (Ocean Health Corporation, Urasoe city) for assistance in Okinawa mosquito collection; David Armitage (OIST) for lab support in Okinawa; Tatiana Tilley for help with Hi-C libraries preparation; Ximena Bernal for providing *Uranotaenia lowii* samples; Ary Faraji for providing *Sabethes cyaneus* samples for head RNA sequencing; Terence Murphy, Françoise Thibaud-Nissen, Vinita Joardar, and the RefSeq team for gene annotations; Jason Banfelder, Linelle Abueg, and the High Performance Computing Resource Center at the Rockefeller University for IT support; Rockefeller University Genomics Resource Center for RNA and Hi-C libraries sequencing assistance; Tatiana Gandlin for mosquito illustrations; the European Synchrotron Radiation Facility (ESRF; Grenoble, France; project number LS-3255) and the Elettra Synchrotron Radiation Facility (Basovizza, Italy; project number 20230106). *Topomyia yanbarensis* and *Malaya genurostris* collections were conducted at the Subtropical Field Science Center (Yona Field), Faculty of Agriculture, University of the Ryukyus, and at Urasoe Daikoen Park in Okinawa Prefecture, with the necessary permissions obtained for each location.

## AUTHOR CONTRIBUTIONS

L.H.-Z. managed the project, obtained and prepared mosquito samples, and performed the majority of genome assembly and data analysis, with additional data acquisition and analyses carried out by co-authors; P.B. conducted the chemoreceptor annotation and performed analysis together with L.H.-Z.; I.A. carried out the collection and analysis of synchrotron data, supervised by P.G. who also participated in synchrotron data collection and analysis; A.E.D. contributed snRNA-seq analysis and assisted with visualization of all figures; J.R.B. prepared the libraries and conducted genome sequencing, supervised by O.F., who also carried out analysis; E.D.J. supervised J.R.B. and O.F. in pipeline and associated analysis approach for high-quality reference genomes assembly; U.P. conducted analysis of viral integrations; N.S. generated the gut blood feeding bulk RNA-seq dataset; T.T. and T.O. together with L.H.-Z. participated in Okinawa field collections, which were led by I.M.; A.M. coordinated the *Sabethes cyaneus* genome assembly generation under the supervision of M.K.N.L.; C.M.S. contributed *Sabethes cyaneus* samples; A.S. contributed *Toxorhynchites rutilus* and *Wyeomyia smithii* samples; L.H.-Z. and L.B.V. together conceived the study, designed the figures, and wrote the paper, with input from all authors.

## EXTENDED DATA FIGURES

**Extended Data Figure 1.**
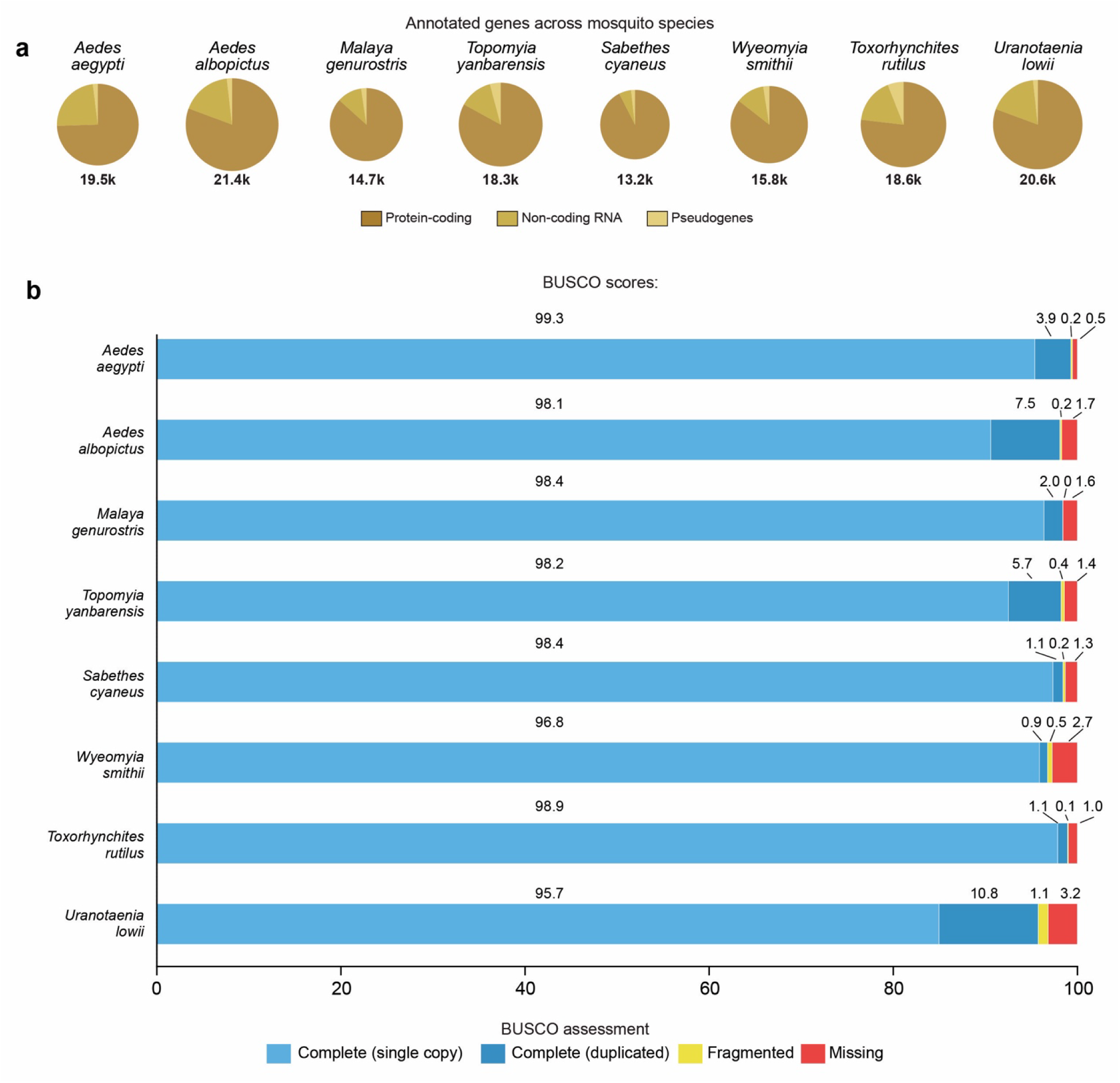
| Gene annotation and assembly completeness across the eight mosquito genomes studied here a, Pie charts of numbers of annotated genes for each of the eight mosquito species partitioned into protein-coding genes, non-coding RNAs, and pseudogenes, with total number of annotated genes below each pie chart. Protein-coding genes exclude TE-derived genes. **b,** BUSCO assessment of genome and annotation completeness for each species, showing the percentage of benchmarking universal single-copy orthologs recovered as complete and single-copy (light blue), complete and duplicated (dark blue), fragmented (yellow) and missing (red). The complete (single-copy+duplicated) percentage per species is indicated in the middle above each bar. Related to Fig. 1d.

**Extended Data Figure 2.**
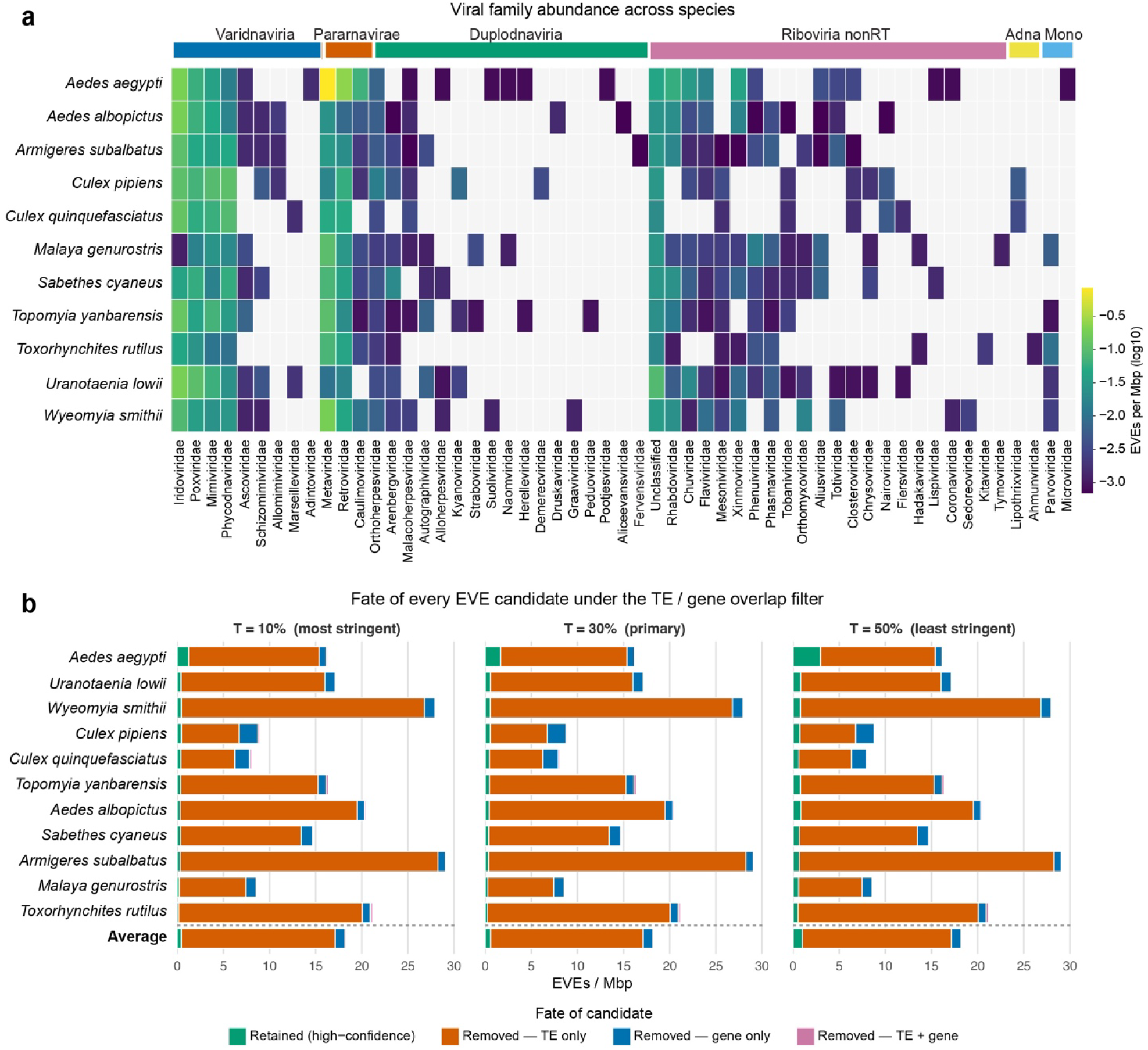
| Endogenous viral element (EVE) content across mosquito genomes. a, Heatmap of EVE abundance (EVEs per Mbp, log10 scale) for each viral family across eleven species (*Aedes aegypti*, the seven species for which genome assemblies were generated here, together with *Armigeres subalbatus*, *Culex pipiens,* and *Culex quinquefasciatus*); gray indicates no EVE detected in that family. **b,** Fate of every candidate EVE under the transposable-element/gene-overlap filtering pipeline, shown at three overlap thresholds (T = 10%, most stringent; T = 30%, primary; T = 50%, least stringent). Stacked bars (EVEs per Mbp) partition candidates by outcome: retained as high-confidence (green), removed for transposable-element overlap only (orange), removed for gene overlap only (blue), or removed for both (pink). Related to Fig. 1.

**Extended Data Figure 3.**
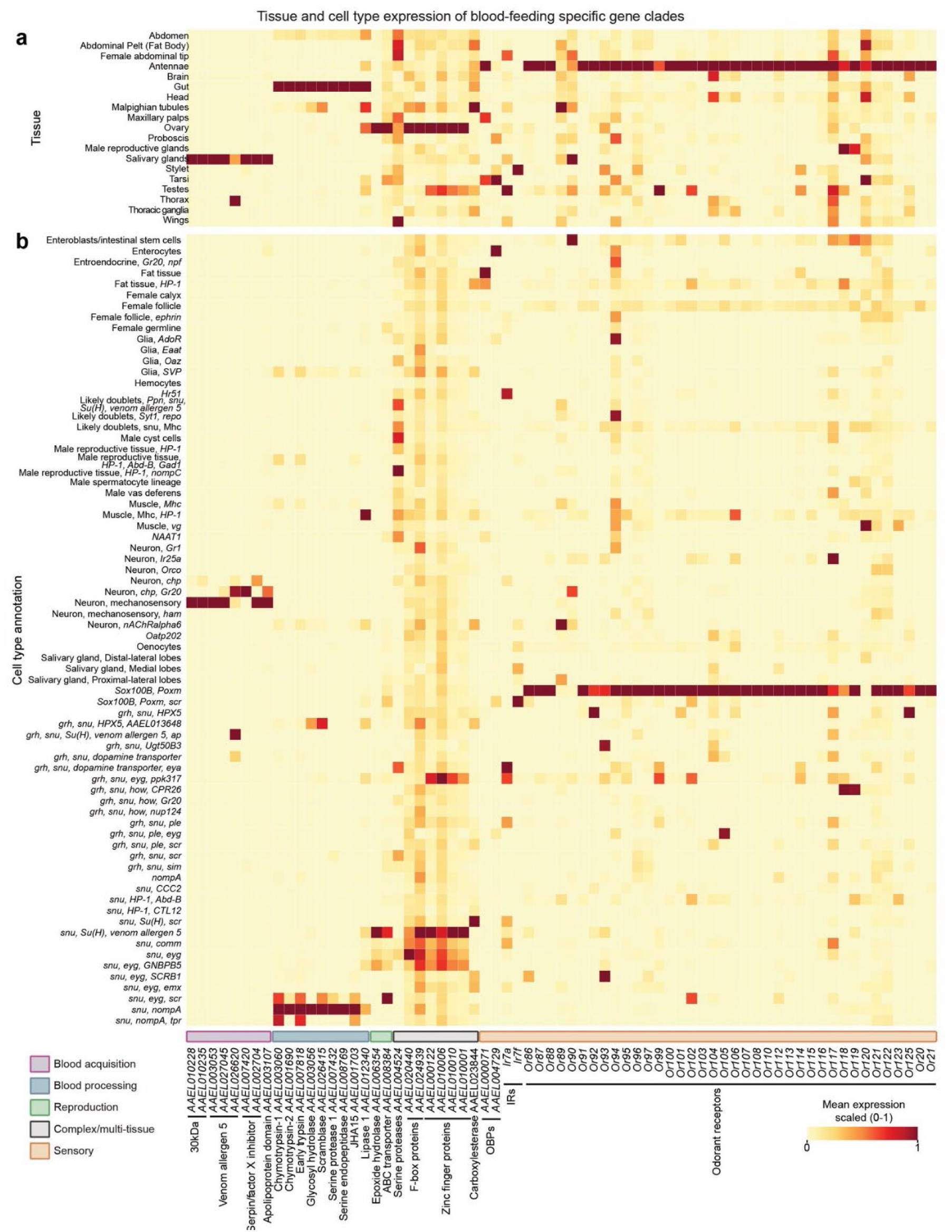
| Tissue– and cell-type expression of 25 blood-feeding-specific gene clades in *Aedes aegypti*. a, Mean expression (scaled 0-1 per gene) of each blood-feeding-specific gene across adult tissues (rows), from single-nucleus RNA-sequencing. Sequencing data from Goldman et al., 2025^49^. **b,** Mean expression (scaled 0-1 per gene) of the same genes across annotated cell types (rows). In both panels, genes (columns) are ordered by five functional categories, with clade names and *Aedes aegypti* gene identifiers given below. Related to Fig. 2c, Fig. 3, and Fig. 4.

**Extended Data Figure 4.**
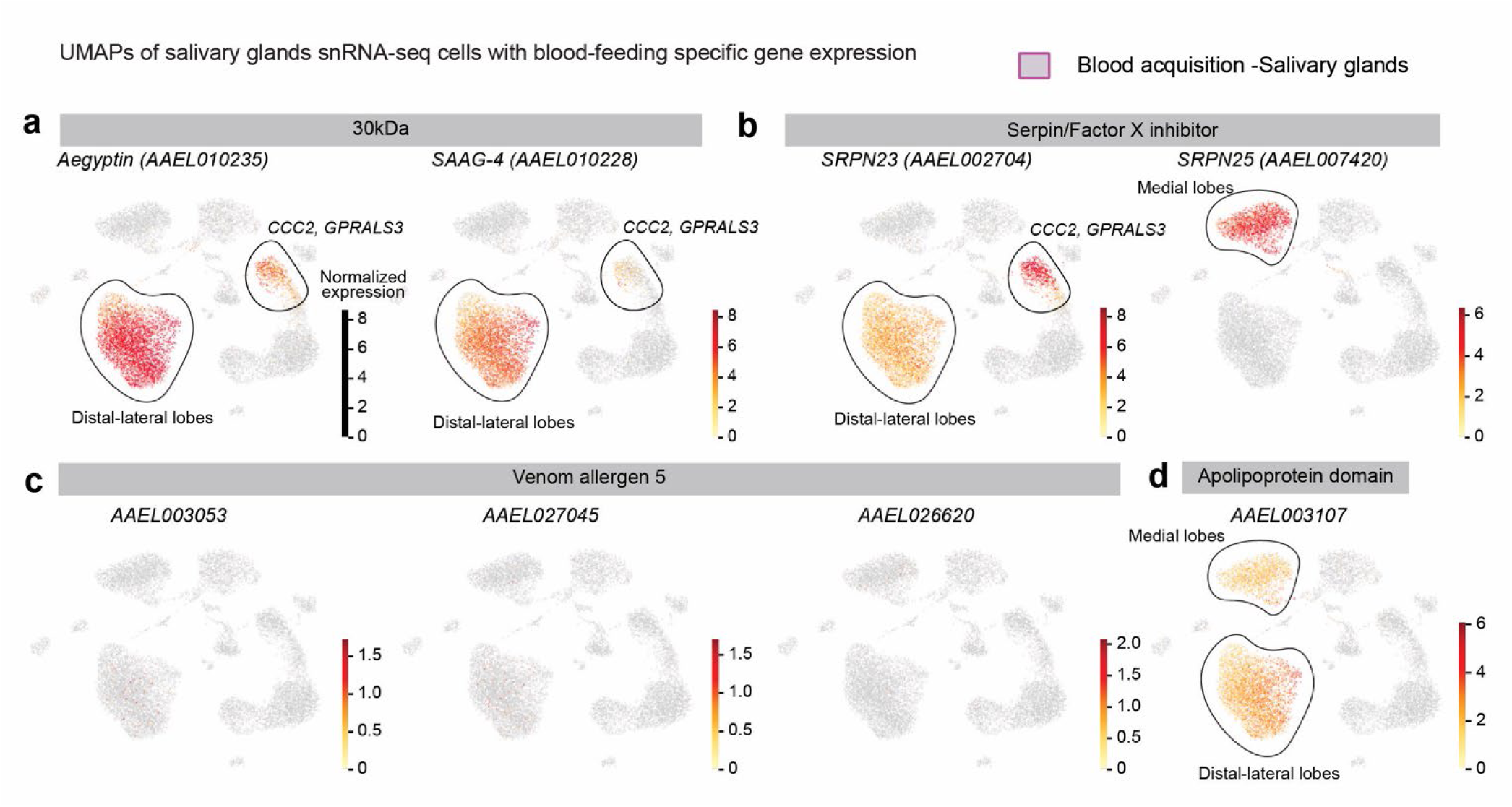
| Single-nucleus expression of blood-acquisition genes across salivary-gland cell types. UMAP projections of *Aedes aegypti* salivary-gland snRNA-seq, each colored by normalized expression of a blood-feeding-specific salivary gene. The distal-lateral and medial lobe cell populations are outlined. Genes shown are **a**, 30-kDa-family genes *Aegyptin* and SAAG-4 expressed in the distal-lateral lobes. **b**, Serpin/Factor-X-inhibitor genes SRPN23 and SRPN25 marking the distal-lateral and medial lobes, respectively. **c**, Venom allergen 5 genes (*AAEL003057* not present in dataset), and **d**, an Apolipoprotein-domain gene expressed in the medial and distal-lateral lobes. Sequencing data from Goldman et al., 2025^49^. Related to Fig. 3c.

**Extended Data Figure 5.**
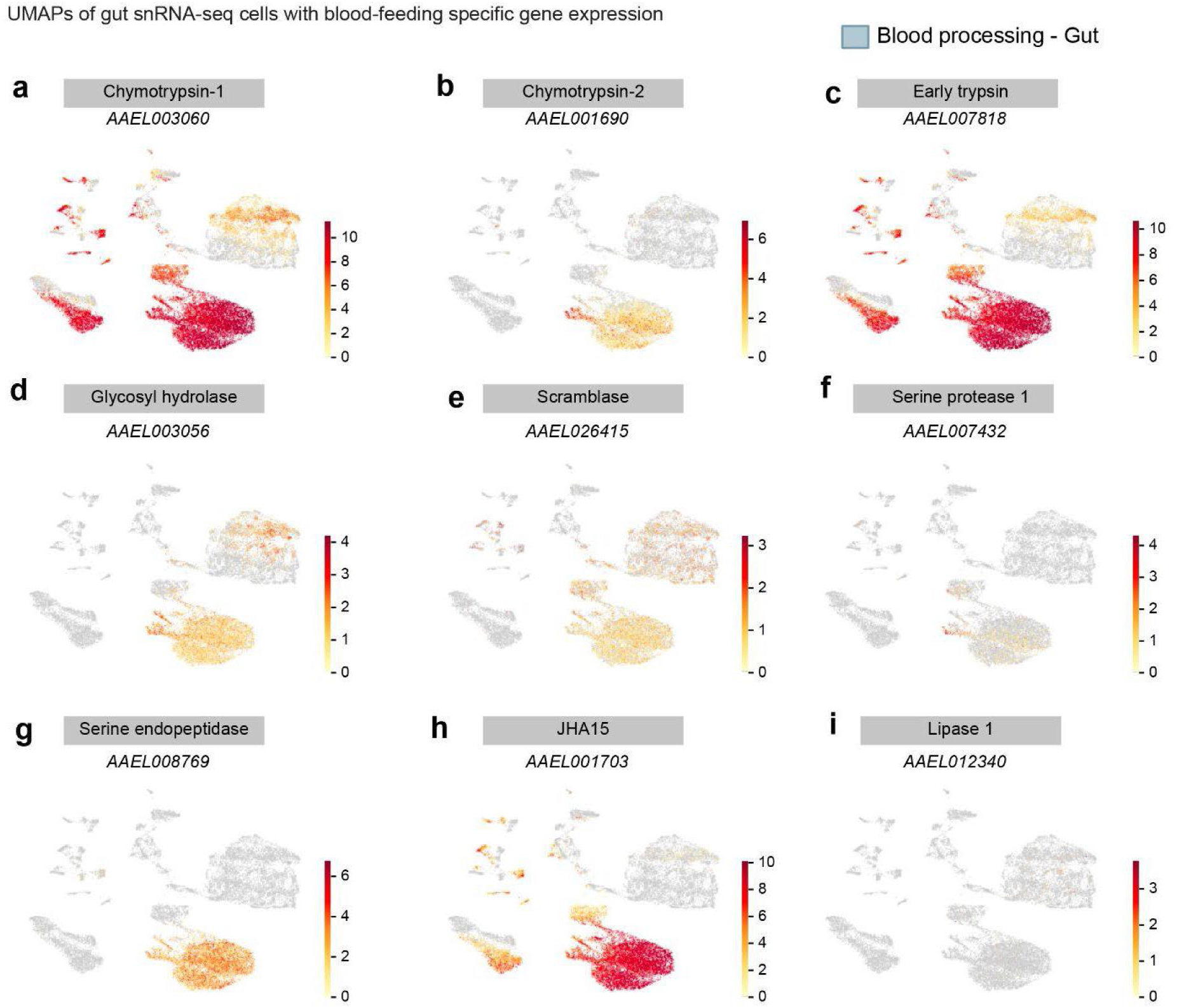
| Single-nucleus expression of blood-processing genes across gut cell types. UMAP projections of *Aedes aegypti* whole-gut snRNA-seq, each colored by normalized expression of a blood-feeding-specific, gut-expressed gene: **a,** Chymotrypsin-1; **b,** Chymotrypsin-2; **c,** Early trypsin; **d,** A glycosyl hydrolase; **e,** A scramblase; **f,** Serine protease 1; **g,** A serine endopeptidase; **h,** JHA15; **i,** Lipase 1. Expression is concentrated in the female gut cells (see male-female cell map in Fig. 3e). Sequencing data from Goldman et al., 2025^49^. Related to Fig. 3d-i.

**Extended Data Figure 6.**
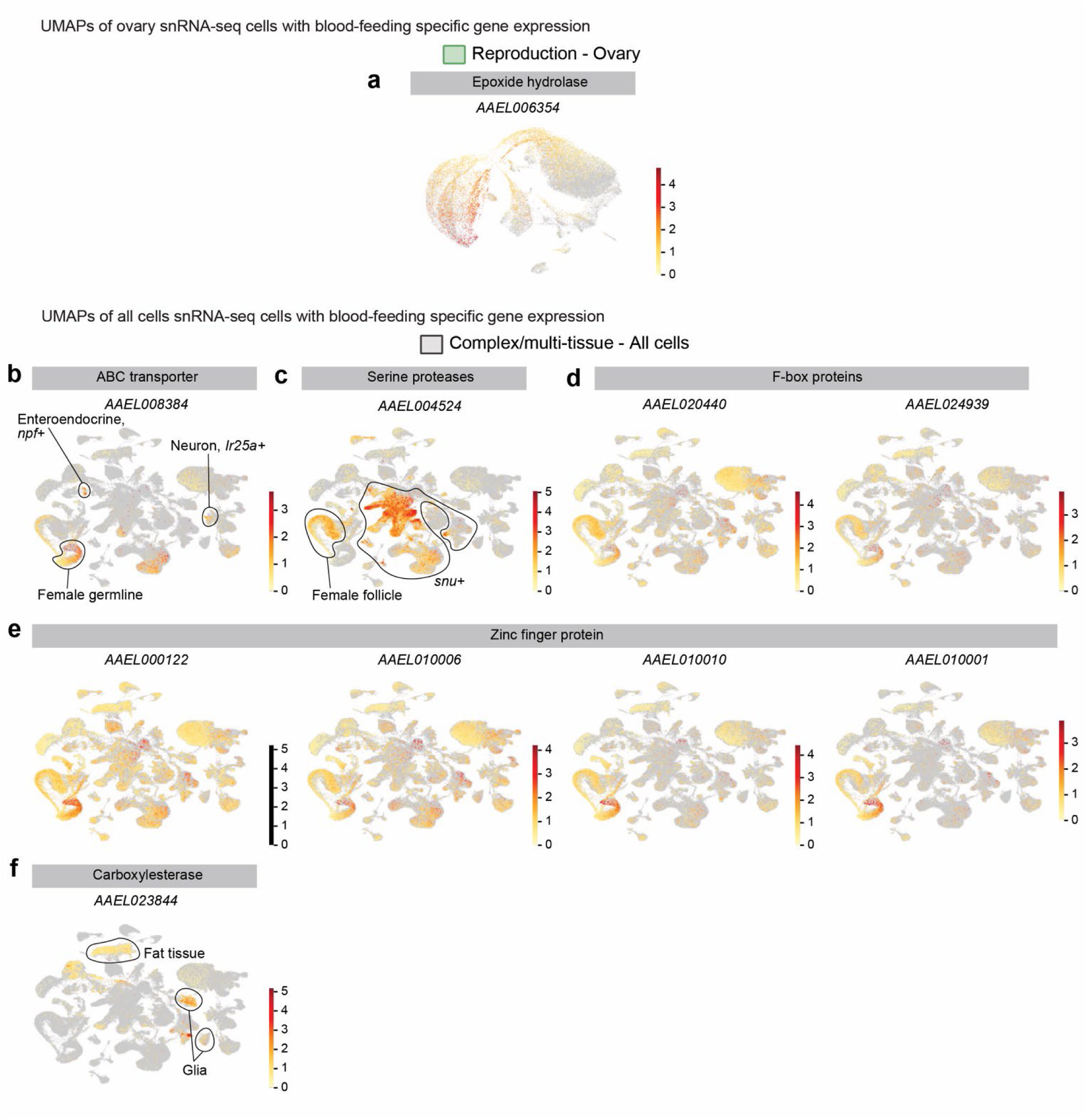
| Single-nucleus expression of reproduction and complex/multi-tissue blood-feeding-specific genes. a, UMAP of *Aedes aegypti* ovary snRNA-seq colored by normalized expression of the reproduction-category epoxide hydrolase. **b-f,** UMAPs of the integrated multi-tissue (“all cells”) atlas colored by normalized expression of complex/multi-tissue genes. Genes shown are an ABC transporter with the enteroendocrine (*npf*+), *Ir25a*+ neuron and female-germline populations indicated (**b**), a CLIP-domain serine protease enriched in the female follicle and in snu+ epithelial cells (**c**), F-box proteins (**d**), Zinc-finger proteins (**e**), and Carboxylesterase CCEae3A, enriched in fat tissue and glia (**f**). Sequencing data from Goldman et al., 2025^49^. Related to Fig. 3j.

**Extended Data Figure 7.**
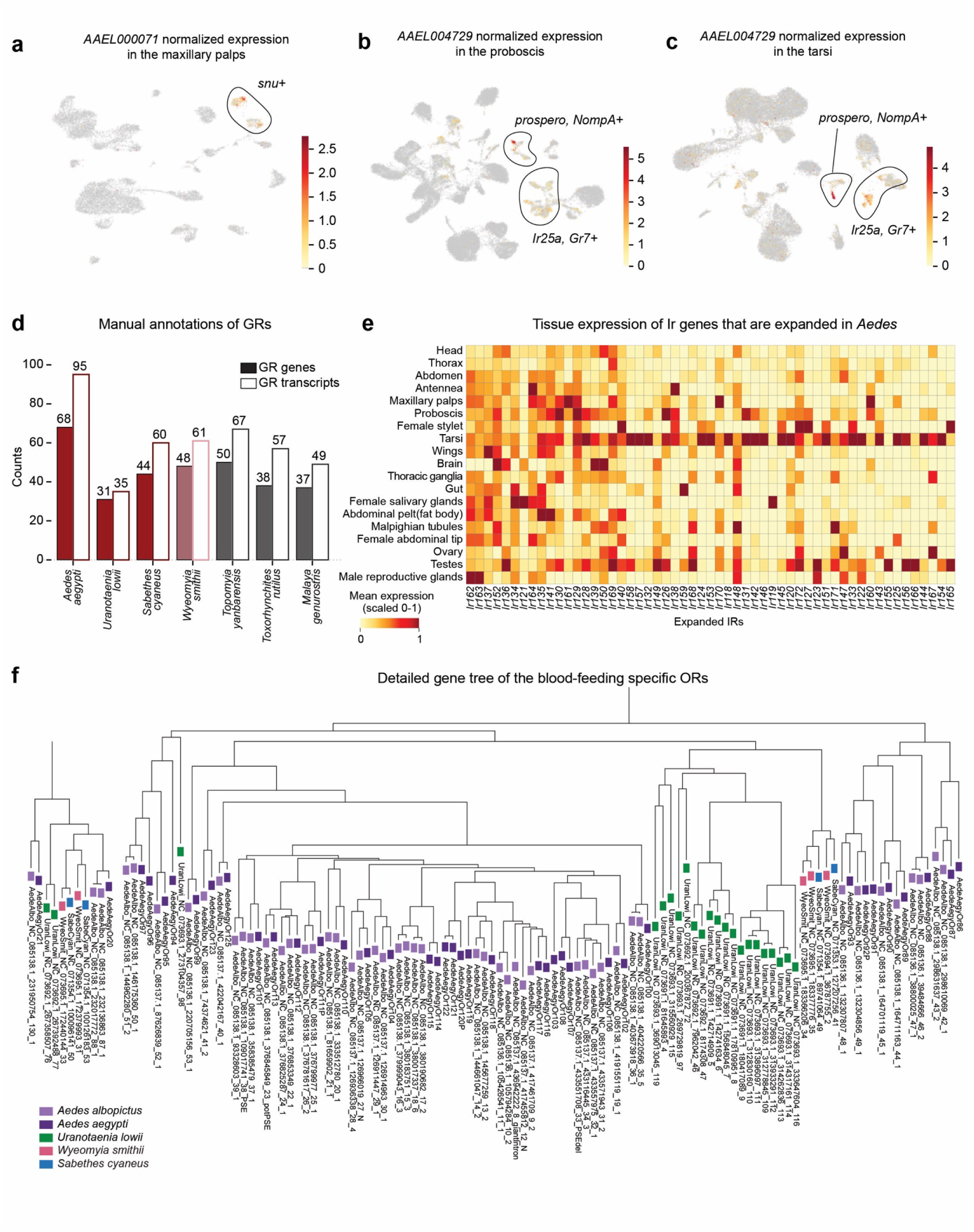
| Chemosensory gene expression and receptor-repertoire annotation. a, UMAP of *Aedes aegypti* maxillary-palp snRNA-seq colored by normalized expression of the blood-feeding-specific odorant-binding protein *AAEL000071*, which marks *snu*+ (epithelial-like) cells. **b,c,** UMAPs colored by normalized expression of the blood-feeding-specific odorant-binding protein *AAEL004729* in the proboscis (**b**) and tarsi (**c**), showing expression in prospero/NompA+ and *Ir25a/Gr7*+ neuronal clusters. **d,** Manually annotated gustatory-receptor (GR) counts per species: number of GR genes (filled bars) and GR transcripts (open bars). *Aedes albopictus* GRs were not annotated (see Methods). **e,** Tissue expression (mean, scaled 0-1) across adult tissues of the ionotropic receptor (IR) genes that are expanded in the *Aedes* lineage. Sequencing data from Goldman et al., 2025^49^. **f,** Detailed maximum-likelihood gene tree of the blood-feeding-specific odorant receptors, with tips colored by species. Related to Fig. 4.

**Extended Data Figure 8.**
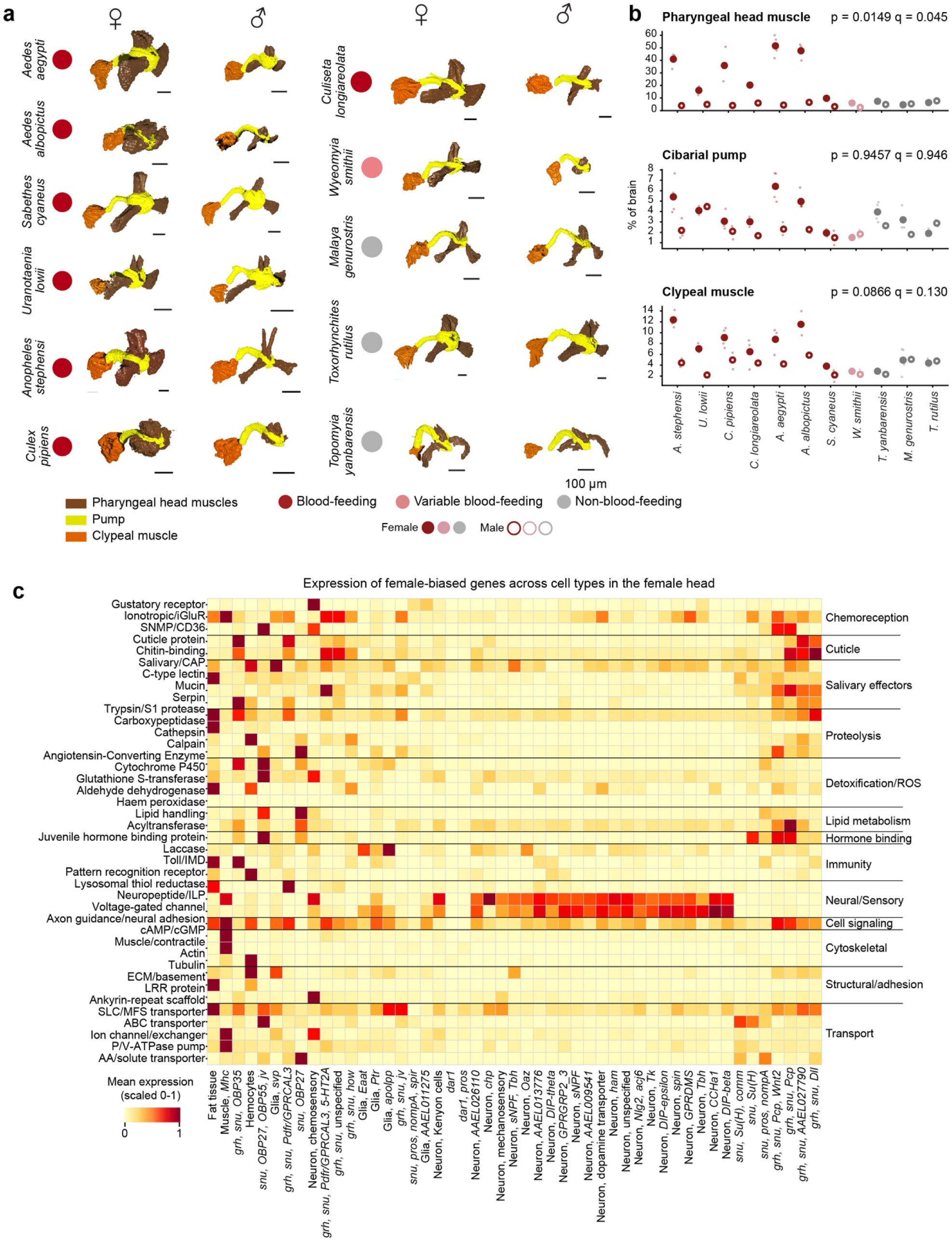
| Sexually dimorphic head musculature and female-biased head gene expression across cell types. a, Volume renderings of the internal head musculature (head muscles, brown; cibarial/pharyngeal pump and associated structures, yellow; clypeal muscles, orange) for females and males of *Aedes aegypti*, the seven mosquito species for which genome assemblies were generated in this study, together with additional control species (*Culiseta longiareolata*, *Anopheles stephensi* and *Culex pipiens*). Species are colored by feeding phenotype. Scale bar, 100 µm. **b,** Relative volume (% of brain) of the head muscle, cibarial pump and clypeal muscle groups in females (filled) and males (open) across species, colored by blood-feeding phenotype. Individual measurements and means are shown. **c,** Mean expression (scaled 0-1) of female-biased head genes grouped by functional gene category across head cell types (columns) in *Aedes aegypti* females. Sequencing data from Goldman et al., 2025^49^. Related to Fig. 5b,d.

**Extended Data Figure 9.**
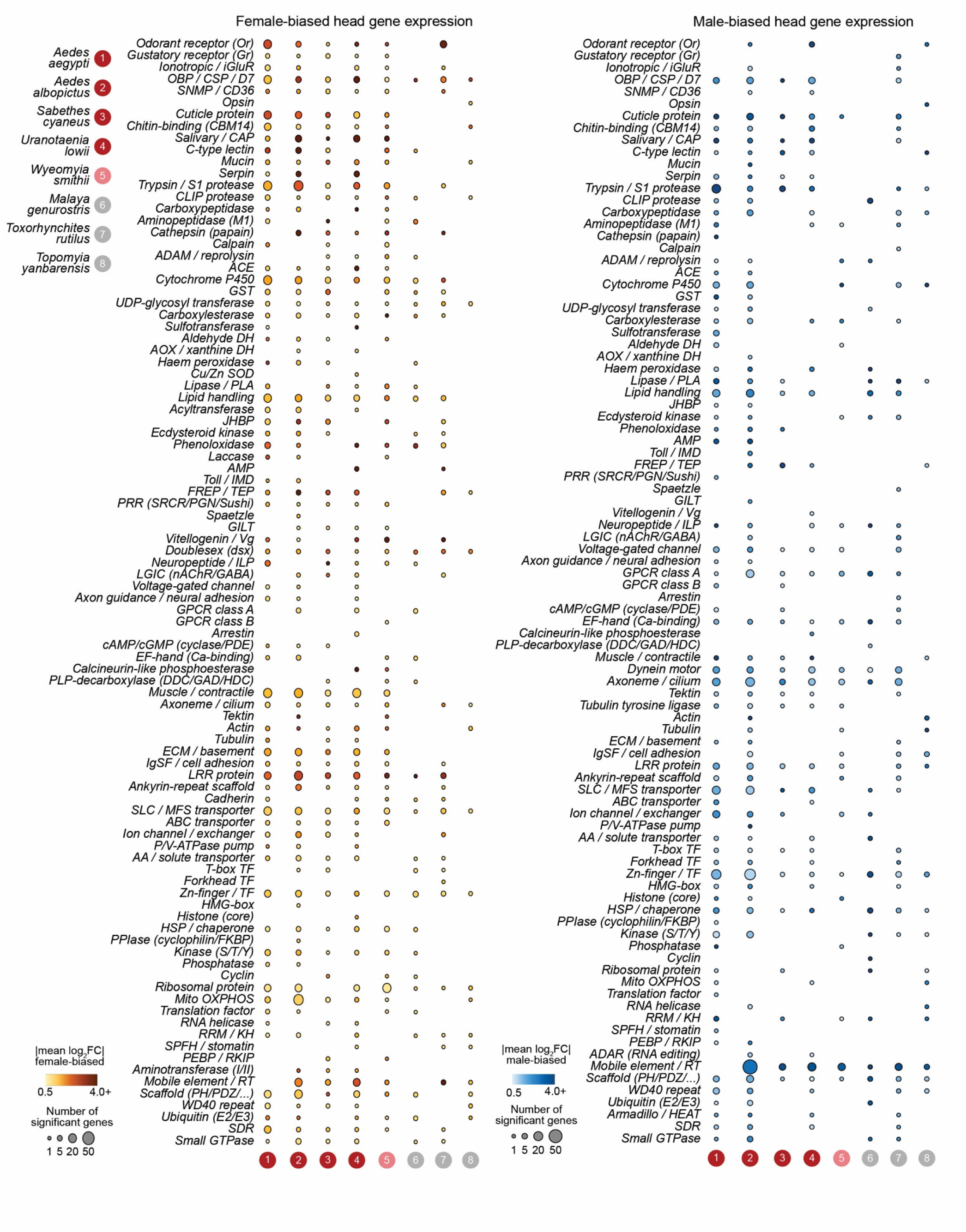
| Functional-category breakdown of sexually dimorphic head gene expression across species. Dot plots summarizing female-biased (left) and male-biased (right) head gene expression by functional gene category (rows) for each of the eight species (numbered 1-8 and colored by feeding phenotype as in Fig. 1). Dot size denotes the number of significantly sex-biased genes in a category and dot color the mean absolute log_2_ fold change. Related to Fig. 5c,d.

**Extended Data Figure 10.**
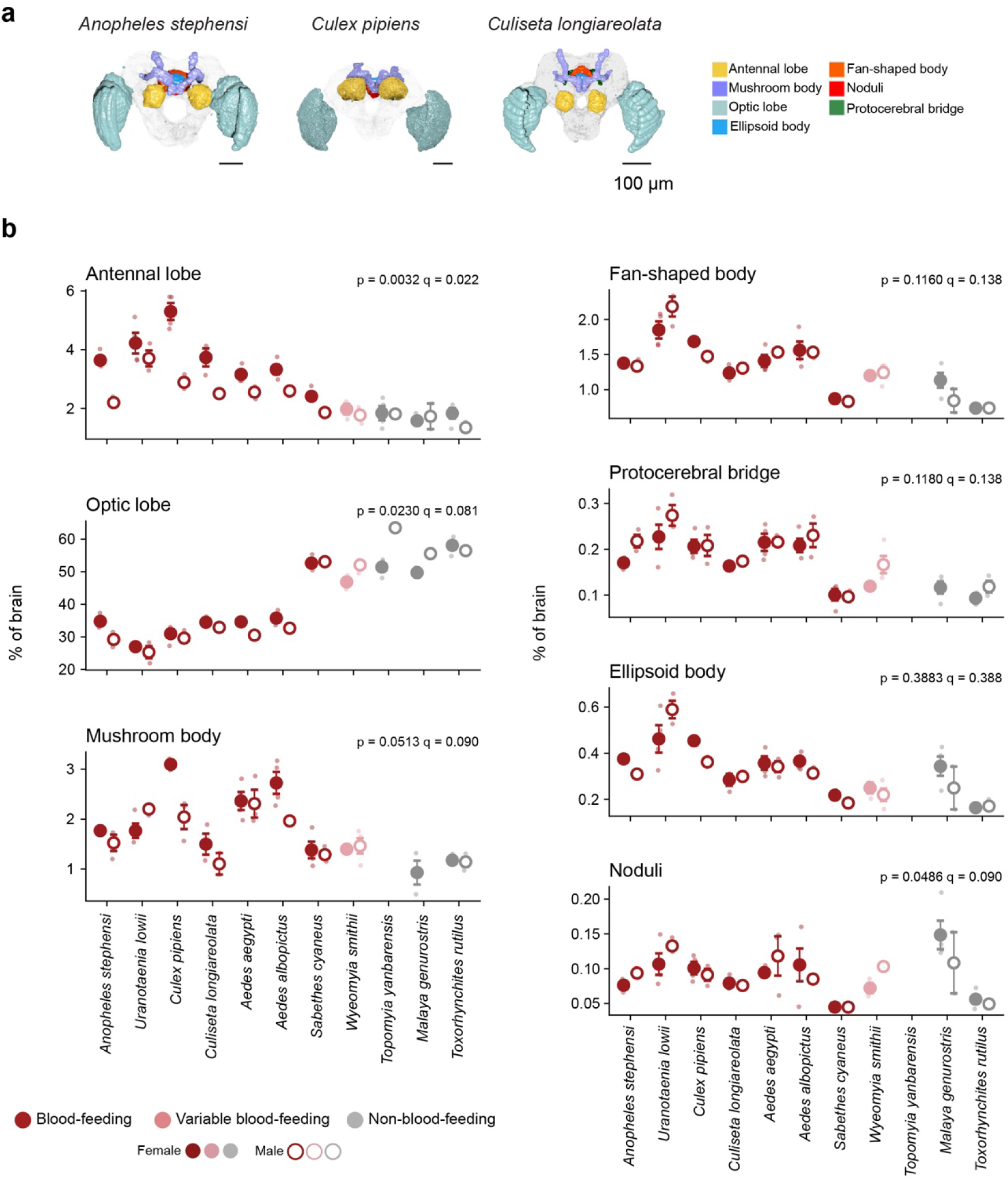
| Whole-brain neuropil reconstruction and regional volumes across species. a, Segmented whole-brain neuropils for three additional species (*Anopheles stephensi*, *Culex pipiens* and *Culiseta longiareolata*), colored by region. Scale bar, 100 µm. **b,** Relative volume (% of brain) of each segmented region in females (filled) and males (open) across species, colored by feeding phenotype. Individual measurements and means are shown. Related to Fig. 5f,g.

## MATERIALS AND METHODS

### Mosquito species counts and taxonomy

Species counts were compiled from the Valid Extant Species of Culicidae checklist maintained by the Mosquito Taxonomic Inventory (Harbach, updated 4 May 2026)^78^. Text was extracted from the PDF checklist and parsed programmatically in Python, and each entry was assigned to its genus and, where given, its subgenus. *Verrallina*, *Nyctomyia,* and *Armigeres* were kept as separate genera rather than folded into *Aedes*, following the Mosquito Taxonomic Inventory’s composite-classification treatment. Species were scored per genus, yielding 42 valid extant genera and 3,729 species. Coloring based on the blood-feeding phenotype was applied at genus level (i.e., the *Wyeomyia* genus is indicated as a blood feeder according to the main genus phenotype).

### Mosquito rearing, strains, and sources

*Rearing at the Vosshall Lab, Rockefeller University* (all species and samples except *Sabethes cyaneus* genome samples): mosquitoes were reared in a constant temperature and humidity environmental room at 26°C with 70 to 80% relative humidity and daily 14-hour light and 10-hour dark cycle (lights on at 7 a.m.).

Data generated in this paper were obtained from the following mosquito strains and sources:

*Aedes aegypti*: Liverpool wild-type laboratory strain (LVP) was initially collected in west Africa and has been maintained at the Liverpool School of Tropical Medicine since 1936 and distributed world-wide.

*Aedes albopictus*: FPA strain. Eggs kindly provided by Dr. Laura Duvall at Columbia University (New York, NY, USA). Foshan inbred strain Pavia A (FPA) wild-type laboratory strain. The FPA strain (*Aedes albopictus* wild type) was initially isolated in Foshan, Guangdong, southeastern China in 1981 and has been cultivated in the lab since then. Mosquitoes were reared in the Vosshall Lab until sample collection and kept afterwards. For all applications, samples were flash frozen and kept in – 70°C until processing.

*Uranotaenia lowii*: MFRU-FL strain. Eggs kindly provided by the Ximena Bernal lab at Purdue University (West Lafayette, IN, USA). Colony was established from egg rafts obtained from the USDA-ARS-CMAVE Mosquito and Fly Research Unit (Gainesville, FL, USA). The mosquitoes were shipped to the Vosshall Lab as egg rafts, which were then reared until sample collection. For all applications, samples were flash frozen and kept in –70°C until processing.

*Sabethes cyaneus*: Majé strain. The Majé strain was isolated by Dr. Woodbridge Foster on Isla de Majé, Lago Bayano, in eastern Panama (8.632, –78.633) in 1988 and maintained continuously at The Ohio State University until 2016. From then onwards, *Sabethes cyaneus* were reared by Chris Stone at the University of Illinois at Urbana-Champaign, IL, US (genome) or maintained in the Ary Faraji laboratory at the Salt Lake City Mosquito Abatement District (head RNA-seq datasets). For the genome, to save on shipping costs, ten individual mosquitoes were lightly squished in 100% ethanol and shipped without cold chain to the Wellcome Sanger Institute, which took about one week. Upon receipt, they were stored in –80°C until extraction. For the head RNA-seq extractions, eggs were shipped to the Vosshall Lab and animals were reared until collection.

*Toxorhynchites rutilus septentrionalis*: Harris County Precinct 4 Biological Control Initiative Integrated Population Colony (IPC). Live material was provided by the Harris County Precinct 4 Biological Control Initiative (BCI; Spring, TX, USA), where this obligately non-blood-feeding species was reared for release to augment local wild populations and as means for biological control of blood-feeding mosquito species. The colony supplied (Integrated Population Colony, IPC) was established in February 2019 from two lines founded from wild material collected in tree cavities and artificial containers in Harris County, Texas, in 2014 and 2018, and has been self-maintained since. Larvae were reared at BCI following a published protocol^71^. Mosquito larvae were shipped to the Vosshall Lab in individual containers to avoid death by larval cannibalism together with batch rearing containers. All mosquito larvae were then separated and reared until sample collection. For all applications, samples were flash frozen and kept in –70°C until processing.

*Wyeomyia smithii*: Harris County Precinct 4 Biological Control Initiative colony, founded from a northern (Ithaca, NY) population. Live material was provided by the Harris County Precinct 4 Biological Control Initiative (BCI; Spring, TX, USA). *Wyeomyia smithii* breeds inside the leaf traps of the purple pitcher plant *Sarracenia purpurea.* Populations north of approximately 40°N do not take a blood meal, whereas more southerly populations produce a first batch of eggs without blood and can blood feed to supplement further egg production. Larvae were collected in October 2019 near Ithaca, New York (42.549 N, 76.266 W) by Dr. Parrish C. Brady (University of Texas at Austin) and used to found a colony at the Harris County Precinct 4 Biological Control Initiative (BCI; Spring, TX, USA), which has been maintained since July 2020. Mosquito larvae were shipped to the Vosshall Lab, reared until sample collection, and kept afterwards. For all applications, samples were flash frozen and kept in –70°C until processing.

*Malaya genurostris* and *Topomyia yanbarensis*: *sourced through field collections in Okinawa, Japan.* The larvae of *Topomyia yanbarensis* are usually found alone at water pools in the bamboo internodes with small holes bored by *cerambycid* beetle (*Abryna coenosa Newman*) in Okinawa. Although bamboo internodes with a small hole (concave) are found commonly in the bamboo forest in the northern part of Okinawa, there are few internodes with holes that penetrate through it. It is difficult to collect many *Topomyia yanbarensis* because larvae breed in nature in a short period of time. In place of the *Cerambycid* beetles, we artificially drilled 5 mm diameter holes in 200 internodes on September 9, 2022 and September 23, 2022, and introduced a small amount of water into each hole with a pipette. After that, a total of 179 internodes were recovered on October 20, November 19, and December 13, 2022. A total of 103 fourth-instar larvae were collected.

The larvae of *Malaya genurostris* are typical taro leaf-axil dwellers in Okinawa and although the number of the larval occurrences depends on the amount of rainfall, they are found throughout the year on giant taro plants (Alocasia spp.), which are very commonly seen at mountain side, park and pasture lands. The larval collections were carried out mainly at Urasoe National Park. Two weeks before starting the larval survey, there were many rainy days, and young leaf axils of taro plants were filled with water, and many Malaya larvae were collected.

Both *Topomyia yanbarensis* and *Malaya genurostris* were reared at the laboratory in Okinawa until sample collections. Samples were placed in Nalgene® Cryogenic Vials (Thermo Scientific 5000-0012), and flash frozen in liquid nitrogen inside a dry shipper (Cryo Express Dry Shipper, MiTeGen TW-CX100). Samples were hand carried frozen inside the dry shipper to the Vosshall Lab in the United States with full import permissions. Upon arrival, the samples were transferred to the –70°C freezer and were kept there until downstream processing.

### DNA extraction and genome sequencing

All sequencing and assembly procedures were done at the Rockefeller University Reference Genome Resource Center (also known as the Vertebrate Genome Lab), except for the genome sequencing and assembly of *Sabethes cyaneus*, which were generated at the Wellcome Sanger Institute, using similar protocols. The protocols and pipelines were based on those developed by the Vertebrate Genomes Project (VGP), Earth BioGenome Project^30,31^, and the Sanger Tree of Life^32^ with modifications for DNA extraction from a single insect.

*All species except Sabethes cyaneus*: For each species, a single male mosquito was homogenized using a pellet pestle, and high molecular weight DNA was extracted using the MagAttract HMW DNA Kit (Qiagen Cat. No. 67563) following a modified protocol described by Kingan et al.^33^. Isolated DNA was sheared using the Diagenode Megaruptor 3 (Cat. No. B06010003). 400 ng of sheared DNA was used to prepare a PacBio library with PacBio barcoded adapters (Pacific Biosciences PN 101-628-400) and size-selected with AMPure PB beads (Pacific Biosciences PN 100-265-900) to remove DNA under 5kb. The quantity of library was measured with a Qubit 3 Fluorometer (Qubit dsDNA HS Assay Kit) and insert size was assessed with the Agilent Femto Pulse. *Toxorhynchites rutilus*, *Uranotaenia lowii*, and *Wyeomyia smithii* were sequenced on PacBio 8M SMRT Cells on the Sequel IIe instrument with binding kit 2.2 (PN 101-894-200), sequencing kit 2.0 (PN 101-820-200) and 30h movies with 2h pre-extension, to generate high fidelity (HiFi) 10-20kb long reads. *Malaya genurostris* and *Topomyia yanbarensis* were sequenced with 40h movies and 2h pre-extension and Binding Kit 3.2 (PN 102-194-100).

Due to the small size of the species and low DNA yield, *Uranotaenia lowii*’s DNA extraction and sequencing was done twice and from two different individuals. Data were combined during assembly.

DNA for Hi-C libraries was extracted from 5 additional male individuals per species. Siblings were used whenever possible. Cross linking and library prep was done using Arima 2.0 Hi-C kit prepped with the Arima Library Prep Module. Libraries were then sequenced on Illumina NovaSeq6000 as paired end, 150nt read length.

*Sabethes cyaneus*: The thorax and abdomen of a single female were disrupted by manual grinding with a blue plastic pestle in Qiagen MagAttract lysis buffer and then extracted using the Qiagen MagAttract HMW DNA extraction kit with two minor modifications including halving volumes recommended by the manufacturer due to small sample size and running two elution steps of 100 μl each to increase DNA yield. The quality of the DNA was evaluated using an Agilent FemtoPulse. Low molecular weight DNA was removed using a 0.8x AMPure XP purification. DNA was sheared using a Diagenode Megaruptor 3 to a 12kb fragment size. Sheared DNA was purified using AMPure PB beads with a 1.8X ratio of beads to sample to remove the shorter fragments and concentrate the DNA sample. The concentration and quality of the sheared and purified DNA was assessed using a Nanodrop spectrophotometer and Qubit Fluorometer with the Qubit dsDNA High Sensitivity Assay kit. A sequencing library was prepared using PacBio Express TPK 2.0 with low DNA input. Sequencing was carried out on the Sequel II system with a 24h run time and 2h pre-extension. For Hi-C data generation, a head from a male specimen was used as input material for the Arima V2 Kit according to the manufacturer’s instructions for animal tissue. The sample was processed through Library Preparation using a NEB Next Ultra II DNA Library Prep Kit and sequenced aiming for 100x depth on an Illumina NovaSeq 6000.

### Genome Assembly and Scaffolding

Raw sequencing reads were first trimmed to remove adapter sequences using Cutadapt^79^ (v3.2, 4.0) (all species except *Sabethes cyaneus*). Contig-level assembly was performed from PacBio long-read data using Hifiasm^80^ (v0.14, 0.15, 0.16), generating a primary assembly and an alternate haplotype assembly. False duplications arising from uncollapsed haplotypic sequence were then removed from the primary assembly using purge_dups^81^ (v1.2.3, 1.2.5), with purged sequences reassigned to the alternate haplotype assembly to produce a curated pair of primary and alternate assemblies. The purged primary assembly was subsequently scaffolded using Hi-C proximity ligation data with SALSA2^82^ (v2.2, 2.3) to produce chromosome-scale scaffolds.

### Hi-C Scaffold Curation

*All species except Sabethes cyaneus*: to further refine and validate the SALSA2-scaffolded assembly, the raw Hi-C reads were aligned to the primary assembly and processed using Juicer^83^ (v2.0) to generate contact matrices. The 3D-DNA pipeline^84^ (v180922) was then applied to the scaffolded assembly to automatically correct misjoins and improve contig ordering and orientation based on Hi-C contact signal. The resulting draft was manually reviewed and curated in Juicebox Assembly Tools^85^ (v1.11.08), where Hi-C contact maps were visually inspected to identify and correct residual scaffolding errors (e.g., misjoins, inversions, or translocations), producing the final curated scaffolded genome assembly. Chromosomes were identified in the Hi-C map of the final assembly, and named according to mosquito chromosome naming convention (shortest chromosome = Chr 1, longest = Chr 2, mid-length = Chr 3)^86^.

For *Sabethes cyaneus*: manual curation was performed using gEVAL^87^, HiGlass^88^, and Pretext^89^. The mitochondrial genome was assembled using MitoHiFi^90^, which performs annotation using MitoFinder^91^.

### Assembly Quality Assessment

Assembly quality and completeness were assessed using multiple complementary approaches: standard assembly contiguity statistics (e.g., N50, contig and scaffold values, total length, QV). Base-level accuracy and k-mer completeness were assessed with Merqury^92^, while raw k-mer spectra from the PacBio reads were analyzed using GenomeScope^93^ to estimate genome size, heterozygosity, and repeat content. Genome completeness was evaluated with BUSCO^94^, using the diptera_odb10 lineage dataset.

### Resolving the *Aedes albopictus* assembly

Because the *Aedes albopictus* genome is highly heterozygous and repetitive, standard assembly approaches produced substantial haplotype duplication. To resolve this, aggressive purging of duplicates was applied both during initial contig assembly (Hifiasm) and in a subsequent purging step (purge_dups), including removal of duplications occurring within contigs rather than only at contig boundaries. Remaining duplications were then resolved through manual curation of the Hi-C contact map, based on a characteristic contact signature: a strong diagonal match between two genomic regions in which one region showed contact signal restricted to only one side of the map, consistent with sequencing reads preferentially aligning to a single, more common copy of a duplicated locus. Contigs displaying this pattern were identified and removed as putative duplicates. The manually curated assembly was then assessed by BUSCO completeness scores and total assembly length to evaluate the trade-off between duplication reduction and retention of gene content.

### RNA extraction and sequencing

*Gene annotation* (all species except *Sabethes cyaneus*): Samples were obtained from larvae, pupae, adult male body (head removed), adult female body, adult female heads, adult male heads, and for the lab grown animals, from eggs as well. For each sample type except for heads, one sample was generated to assist with the gene annotation process. Heads were extracted in four independent replicates to examine sexually dimorphic head gene expression (see below) and also assisted in gene annotations. *Sabethes cyaneus gene annotation sample*: RNA was extracted from a whole specimen shipped at room temperature in 100% ethanol. RNA was extracted using TRIzol, according to the manufacturer’s instructions. An RNA library was created using the directional NEB Ultra II stranded kit and sequencing was performed on an Illumina HiSeq 4000.

*Head sexually dimorphic gene expression*: the mosquitoes were first cold anesthetized by placing them on ice. Using dissection forceps, the animals’ heads were gently removed and collected into sterile 500 µl Eppendorf tubes. 4-5 males and 4-5 females (consistent per species) were collected per species, in 4 separate replicates.

RNA extraction was performed using the PicoPure Kit (Thermo Fisher, KIT0204) with the following modification for homogenizing tissue: instead of lysis buffer, 100 µL of TRIzol (Thermo Fisher, 15596026) was added to the collection tube on ice. Tissues were manually homogenized using a Pellet Pestle Motor (Kimble, 749540) and an RNase-Free pellet pestle (VWR, KT749510-0590) for 30 s. An additional 140 µL was added, bringing the final volume to 240 µL. Tubes were left at room temperature for 5 min before 48 µL of chloroform:isoamyl alcohol 24:1 (Sigma, C0549) was added. Tubes were hand-shaken for 30 s and left to stand for 2 min before centrifuging at 12,000 RPM for 15 min at 4 °C. The aqueous TRIzol layer was then removed and added into the PicoPure column, up to 130 µL at one time. Subsequent steps were performed according to PicoPure manufacturer’s instructions, including DNase treatment. Extracted RNA was then used as a starting material to generate RNA-seq libraries using Illumina TruSeq stranded mRNA LT kit (Illumina, 20020594), following the manufacturer’s protocol. Libraries prepared with unique dual indexes were pooled at equal molar ratios. Sequencing was performed at The Rockefeller University Genomics Resource Center on the Illumina NovaSeq 6000 sequencer to generate 150 bp paired-end reads, following the manufacturer’s protocol.

### Gene annotation

Assembly and supporting RNA-seq samples were submitted to the NCBI RefSeq Eukaryotic Genome Annotation Pipeline for annotation^95^. RefSeq’s accession numbers per species are detailed below (under “Data availability”).

### Identification and annotation of transposable and repetitive elements

Repeat and transposable elements were identified *de novo* in each genome assembly using RepeatModeler^96^ (v2.0.4) with the NCBI/RMBlast search engine (-engine ncbi) and the LTR structural discovery pipeline enabled (-LTRStruct). All other parameters were left at their defaults. The species-specific repeat library produced by RepeatModeler (-families.fa) was then used to quantify, classify and mask repetitive and transposable elements in the corresponding genome FASTA using RepeatMasker^97^ (v4.1.5) with the RMBlast search engine (-e rmblast) and otherwise default settings. The resulting tables and annotations are available in Extended Data^36^.

### Analysis of viral integrations

A database of viral proteins from 12,493 species (NCBI Virus, Jan 2025) was compiled and partitioned into six ICTV-based groups: Adnaviria, Duplodnaviria, Monodnaviria, Varidnaviria, Pararnavirae, and non-reverse-transcribing Riboviria. Candidate EVEs were identified using EEfinder^98^ as an initial similarity screen, then processed with a custom pipeline (EVE-classifier) for de-duplication, re-validation, taxonomic assignment, and genomic-context annotation. The EEfinder screen and the custom EVE-classifier pipeline is archived in Extended Data^36^.

Redundant candidate EVEs were reduced in two steps: overlapping hits at the same locus were collapsed to the single longest EVE, and remaining EVEs within 30 bp sharing the same viral genus were merged. Each resulting locus was re-validated and assigned to a single viral protein via BLASTX (best hit by bitscore, ties broken by E-value), yielding one group/family/genus/species assignment per EVE. Unclassified hits were retained but excluded from family-diversity counts.

To annotate genomic context, EEfinder’s reverse-similarity filter was omitted (it can discard true, host-like EVEs) and replaced with intersection against host gene/TE annotations. Each EVE was classified by overlap with exons (characterized genes only), introns, intergenic space, and annotated TEs. A high-confidence EVE set was defined as loci with ≤30% overlap with TEs or characterized exons (tested also at 10%/50%; 30% used as primary threshold), since TE overlap reflects genomic position rather than sequence identity and was not treated as an evidentiary signal. Per-species and per-family summaries (EVE counts, lengths, taxonomic diversity, genomic context, overlap fractions) were generated, normalizing burden by genome size (density and occupancy). Context/overlap percentages were computed from summed counts rather than averaged across groups to avoid bias from small groups, then merged into cross-species matrices and a family-prevalence table.

### InterProScan

The mosquito protein-coding sequences (the RefSeq protein FASTA file of each assembly, GCF_*_protein.faa) were used as input for domain analysis using InterProScan v5.63-95.0^99^. InterProScan was run with default settings, i.e., against the default set of member databases with no restriction of the analyses performed (no –appl flag) and with no additional output options specified. Domain annotations output per species is available in Extended Data^36^.

### Filtering Transposable Element (TE)-derived genes

Initial runs of orthology assignment (see below) revealed high prevalence of genes that are annotated as protein coding and bear strong characteristics of intact transposable-elements derived genes (for example, fully duplicated, containing only transposable domains). To focus our orthology assessment on genes that support endogenous functions across the mosquitoes, we developed a semi-manual procedure to omit such TE-derived genes from our analysis. To this end, we used InterProScan output to flag genes which contained TE-related domains (key words: transpos*, RT_L, RT_N, reverse transcriptase, integrase, Gypsy, DUF5641, DUF1759, retro). The flagged genes were then manually inspected to determine whether they contain additional non-TE domains – in which case they were kept in the analysis – or contain only TE elements, in which case they were not included in the downstream orthology analysis. Full lists of TE-like genes are available in Extended Data^36^. The filtered protein-coding gene set per species was then used for downstream processing for all applications.

### Synteny analysis

The global synteny map was created using GENESPACE^100^ (v1.3.1) run with default parameters.

### Identifying gene clades across species (orthology analysis)

OrthoFinder3^40^ (v3.0.1b1) was run on the protein sets of the following 15 genome assemblies: *Aedes aegypti (*GCF_002204515.2), *Aedes albopictus (*GCF_035046485.1), *Malaya genurostris* (GCF_030247185.1), *Sabethes cyaneus* (GCF_943734655.1), *Toxorhynchites rutilus* (GCF_029784135.1), *Topomyia yanbarensis* (GCF_030247195.1), *Uranotaenia lowii (*GCF_029784155.1), *Wyeomyia smithii* (GCF_029784165.1), *Culex pipiens* (GCF_016801865.2), *Culex quinquefasciatus* (GCF_015732765.1), *Armigeres subalbatus* (GCF_024139115.2), *Anopheles gambiae* (GCF_943734735.2), *Anopheles coluzzii* (GCF_943734685.1), *Anopheles darlingi* (GCF_943734745.1) and *Culicoides brevitarsis* (GCF_036172545.1). The input protein fasta files were collapsed to retain only the longest isoform per gene and filtered for TE-derived genes as described above.

OrthoFinder was run with the multiple-sequence-alignment gene-tree inference method (-M msa) and otherwise default parameters. For downstream processing, we used the phylogenetic node that was defined by the inclusion of all culicine species and without the *Anopheles* and non-mosquito outgroup *Culicoides brevitarsis* (N3) and retrieved its hierarchical orthogroups output.

### Species phylogeny

The species phylogeny and the cladograms shown throughout the paper are based on the species tree inferred by OrthoFinder from the orthogroups described above. See Extended Data^36^ for tree file. For the neuroanatomical analyses, which included additional species and required a time-calibrated tree for the phylogenetic generalized least squares models, the time-calibrated mosquito phylogeny of Soghigian et al.^19^ was used instead.

### Identification and manual validation of gene loss

To identify blood-feeding-specific genes and their convergent loss, we extracted Hierarchical Orthogroups (HOGs) in which all blood feeding species had at least one gene copy, while all the non-blood feeders had no matching orthologs identified. The variable blood feeder, *Wyeomyia smithii*, was held out of this analysis and examined later for whether it had orthologs to the identified orthogroups.

Manual validation of loss was done using the following procedures:

*Protein-to-genome alignment.* The protein sequences of the candidate lost gene clades, taken from *Sabethes cyaneus*, were aligned back to the genome assemblies of the non-blood-feeding species and the variable blood feeder (non-blood feeders: *Malaya genurostris*, *Toxorhynchites rutilus* and *Topomyia yanbarensis,* variable blood feeder: *Wyeomyia smithii*) using miniprot^101^ with the command miniprot –t16 –-gff –I –u –p 0

“SPECIES“.mpi “LOST_PROTEINS”.fa. The resulting GFF alignments were intersected with the RefSeq annotation of the corresponding genome using bedtools^102^ intersect with –wa –wb –loj, to identify alignments overlapping with annotated features. The alignment outputs and overlapping features were then manually inspected to identify mis-annotated regions, genes that were wrongly excluded, and any pattern that would indicate the presence of the gene in the non-blood-feeding species.

*Microsynteny analysis*. For each candidate lost gene clade, microsynteny was evaluated using *Sabethes cyaneus* as the reference species. For every reference *Sabethes* gene, the 10 annotated genes immediately upstream and the 10 immediately downstream on the same scaffold were taken as syntenic anchors. Each anchor was mapped to its hierarchical orthogroup (node N3, see above) and, for each of the other 10 species, back to the orthologous gene or genes in that species’ RefSeq annotation along with its genomic position. The anchors “map” was then manually inspected to identify regions in which syntenic orthologs are retained. These anchors were then used to define the putative loss region in the non-blood feeders for a targeted downstream analysis across the species.

*tblastn around loss regions*. Candidate gene losses in the three non-blood-feeding species (*Topomyia yanbarensis*, *Malaya genurostris*, and *Toxorhynchites rutilus*) and the variable blood feeder (*Wyeomyia smithii*) were validated at the sequence level by tblastn (NCBI BLAST+). For each lost gene candidate, the protein sequence of its *Sabethes cyaneus* ortholog was used as the query. The putative genomic interval that contained the lost gene (based on the previously described anchors) at each locus was expanded by 2.5Mb on either side. The corresponding genomic window was extracted from the target assembly (samtools faidx) and compiled into a per-window nucleotide BLAST database. tblastn was run with an e-value threshold of 1 x 10^−3^, word size 3, and a maximum of 50 target sequences. High-scoring segment pairs were re-projected onto scaffold coordinates and intersected with the putative loss interval and with annotated gene models for manual inspection of the putative loss regions.

Manual inspection was performed by examining each of the “loss” windows across the species for unannotated genes, pseudogenes, or orthogroups that only correspond to the three non-blood-feeders (and therefore represent non-blood-feeding-specific sequence diversification rather than loss). The results of this validation excluded two genes that were missing annotations in the non-blood-feeding species but were present in the genome, reducing the initially identified 26 blood-feeding-specific gene clades to 24. An additional gene clade was then added by manual inspection of the chemoreceptor gene trees and the chemoreceptor orthogroups were corrected according to the trees: for the rapidly evolving chemoreceptor families, in which not all genes were annotated by RefSeq’s pipeline, gene losses were further inspected by using the construction of full gene trees and searching for branches that contain all and only blood-feeding species genes (see below).

### Categorizing lost gene clades

The lost gene clades were categorized into five broad categories: 1) Blood acquisition, defined by salivary gland expression and functions. 2) Blood processing, defined by gut expression and functions. 3) Reproduction, defined by ovary-specific expression. 4) Complex/multi-tissue, defined as genes that were expressed across multiple tissues and not confined to a specific putative function. 5) Sensory, defined as genes of chemosensory families that are expressed in main sensory tissues (i.e., antenna, tarsi, maxillary palps, proboscis, stylet).

Each lost gene clade was manually inspected to determine its domain annotation using InterProScan output, define its expression pattern in *Aedes aegypti* using the Mosquito Cell Atlas^49^ across all tissues and cell types, and searched against existing literature across species. Genes that had no obvious expression in sugar-fed animals, but had strong expression in a specific tissue at post blood-feeding timepoint(s) were assigned to a function based on the induced expression (e.g., Lipase-1).

### Single nucleus RNA-Seq data and analysis

All datasets used are publicly available in Extended Data^36^ and analysis scripts were adapted from the previously published analyses^103^. For UMAP and heatmap visualizations, normalized expression was calculated as ln((raw count / total cell count) x median total counts across cells + 1) as described before^49^. Heatmaps categorized by tissue or by annotation display mean normalized expression per gene, scaled 0-1 based on each gene’s minimum and maximum value using Scanpy’s sc.pl.matrixplot (standard_scale=’var’), with visualization via seaborn’s sns.heatmap.

The grouped heatmap comparing lost and retained odorant receptors plots unscaled mean normalized expression per gene within each OR cluster, with non-OR clusters excluded. The “Rbfox1 (chemoreceptor unknown)” cluster was renamed “Or20” for this figure, and clusters are ordered left to right from highest to lowest value. UMAP visualizations of normalized expression were generated using sc.pl.umap. Panels showing cell sex plot the recorded sex of each cell directly, with no calculated value. Figure 4h displays the summed normalized expression of all genes in the lost-OR group per cell, with the color scale capped at a normalized expression value of 6 and zero values shown in grey.

### Analysis of sex-bias in the neuronal clusters in the antenna

Cell-composition sex bias in antennal chemosensory neuron clusters was quantified from single-nucleus RNA-seq nucleus counts, using per-cluster female and male counts (assigned by the sex of the sample of origin) across receptor-defined clusters. Because the data comprised four female samples and one male sample, female counts were divided by four to normalize to a 1:1 female-to-male sampling ratio, and a normalized female fraction was computed per cluster as the corrected female count divided by the sum of the corrected female and male counts.

### Mosquito rearing and blood feeding for midgut bulk RNA-sequencing

*Aedes aegypti* wild-type (Liverpool) mosquitoes were reared in an environmental chamber maintained at 26°C ± 2°C with 70-80% humidity, with a photoperiod of 14h light: 10 h dark as previously described^67^. Embryos were hatched in 1L hatching broth: one tablet of powdered Tetramin (TetraMin Tropical Tablets 16110M) in 1L of deionized water, then autoclaved. Larvae were reared in deionized water (3 L total) and fed 2 crushed Tetramin tablets on the first day post-hatching and 2 tablets daily thereafter. To maintain a low rearing density, approximately 400 larvae were kept in 3 L of deionized water from the L3 to L4 stage. Adult mosquitoes were supplied with unlimited access to 10% sucrose solution (w/v in deionized water), delivered in a glass bottle (Fisher Scientific FB02911944) with a cotton dental wick (Richmond Dental 201205), and were kept in 30 × 30 × 30 cm BugDorm-1 Insect Rearing Cages (BugDorm DP1000). Males and females were housed together in these cages from eclosion, allowing females to mate freely.

For all blood-feeding time points, 13-15-day-old mated females were fed defibrinated sheep blood (Hemostat Laboratories, DSB100) supplemented with 2mM adenosine 5’-triphosphate (ATP) (Sigma-Aldrich, A6419) in a 25 mM aqueous sodium bicarbonate buffer using a “blood puck” artificial membrane as described before^55^.

After feeding, females that were not fed or partially engorged were removed. Fully engorged females were kept in their original rearing conditions with continuous access to 10% sucrose until dissection (3h, 6h, 12h, 24h, 48h or 72h post blood meal).

### Bulk RNA-sequencing of mosquito midgut

Adult wild-type *Aedes aegypti* (Liverpool) mosquitoes aged 14-16 days, that were sugar fed (for non-blood-feeding control or males) or blood fed (3h, 6h, 12h, 24h, 48h or 72h prior) were aspirated using an oral aspirator (John W. Hock Company 612) into a 16-ounce container (Webstaurant KH16A-J8000) and were sealed using double 0.8mm polyester mosquito netting (ahh.biz F03A-PONO-MOSQ-M008-WT), then anesthetized on ice for up to 30 min, or until dissections were complete. Midguts were dissected on ice in ice-cold RNase-free 1X phosphate-buffered saline (PBS) (Invitrogen, AM9625). They were moved using forceps into 0.5mL Eppendorf LoBind microcentrifuge tubes (Sigma-Aldrich, Z666521), and immediately snap-frozen on a cold block (Simport, S700-14) pre-chilled to –78°C on dry ice. Extreme caution was taken during tissue dissection to prevent contamination from other mosquito tissues. Each dish and each pair of forceps were carefully cleaned with 70% ethanol and RNase-away (Thermo Fisher, 7003) after each dissection. All replicates for each experimental group were dissected in parallel to avoid artifacts and batch effects. Dissected tissue was stored at –80°C for up to 1 week until RNA extraction. Three midguts were used for each female replicate, 25 midguts per male replicate, and 5 replicates were prepared per experimental group.

RNA extraction was performed as described above.

Samples were run on a Bioanalyzer RNA Pico Chip (Agilent, 5067–1513) to determine RNA quantity and quality. RNA quantity was re-verified with a Qubit 2.0 Fluorometer using the RNA HS Assay Kit (Invitrogen, Q32855). The three biological replicates with the most consistent RNA yield and RNA integrity across conditions were then used for library preparation and sequencing.

Of total RNA, 100 ng was used for library preparation for all samples, except male samples with lower yield, for which a minimum of 25 ng was used due to the smaller male midgut size to generate RNA-seq libraries using the Illumina TruSeq stranded mRNA LT kit (Illumina, 20020594), following the manufacturer’s protocol. Libraries prepared with unique dual indexes were pooled at equal molar ratios. Sequencing was performed at The Rockefeller University Genomics Resource Center on the Illumina NovaSeq 6000 sequencer using V1.5 reagents, the SP flow cell, and NovaSeq Control Software V1.7.0 to generate 150 bp paired-end reads, following the manufacturer’s protocol. Data were demultiplexed and delivered as FASTQ files for each library. Sequencing reads have been deposited at the National Center for Biotechnology Information (NCBI) Sequence Read Archive (SRA) under BioProject ID PRJNA1502948.

### Analysis of midgut RNA-seq data and publicly available RNA-seq datasets

Our dataset, along with the publicly available anterior and posterior midgut dataset^48^ and ovary dataset^55^ were processed uniformly through the same pipeline: adapter and quality trimming with Cutadapt, transcript quantification with kallisto, and gene-level summarization and normalization with tximport and DESeq2, as described below. Each dataset was processed and normalized independently.

Read quality was assessed with FastQC^104^. Paired-end reads were then trimmed with Cutadapt^79^ (v5.2) using the following Illumina TruSeq adapter sequences:

AGATCGGAAGAGCACACGTCTGAACTCCAGTCA (read 1, –a) AGATCGGAAGAGCGTCGTGTAGGGAAAGAGTGT (read 2, –A)

We used a quality cutoff of 20 (-q 20) and a minimum retained read length of 36 nt (-m 36).

Bulk RNA-seq reads from *Aedes aegypti* midgut (female blood-meal time course: non-blood-fed and 3, 6, 12, 24, 48 and 72 h post-blood-meal, plus male midgut, three biological replicates per condition) and ovary (non-blood-fed and post-blood-meal timepoints) were quantified against the *Aedes aegypti* reference transcriptome (“Aaeg_VBJo19”, see Extended Data^36^) using kallisto^105^ (v0.52.0). kallisto quant was run in paired-end mode with 100 bootstrap replicates (-b 100).

A transcript-to-gene map was constructed from the *Aedes aegypti* annotation (AaegLVP_VB58-Jove19.gff^103^) by taking every feature of type mRNA, lnc_RNA, ncRNA, tRNA, rRNA, snRNA, snoRNA, miRNA, pseudogenic_transcript or transcript and pairing its ID attribute (transcript) with its Parent attribute (gene). Transcript-level abundances were imported and summarized to gene level in R with tximport^106^ (type = “kallisto”, ignoreAfterBar = TRUE, ignoreTxVersion = TRUE). Gene-level counts were loaded into DESeq2^107^ via DESeqDataSetFromTximport with a design of ∼ 1, and size factors were estimated using the median-of-ratios method to produce normalized counts. Technical replicates sequenced across two lanes were summed per biological replicate prior to plotting.

### Transcriptional ranking of blood-processing genes

All genes were ranked within each sample by their DESeq2-normalized read count mean across replicates, from the most to the least abundant, and the rank of each blood-processing gene clade is shown (rank 1 = the most highly expressed gene).

### Marker-gene statistics from the single-nucleus data

Marker-gene statistics reported for the single-nucleus data (log fold change and adjusted p-value) were retrieved from the UCSC Cell Browser session hosting the published *Aedes aegypti* single-nucleus atlas^49^.

### Manual annotation of chemoreceptors

Chemosensory receptor genes of three gene families (ORs, IRs, GRs) were annotated in the newly sequenced genome assemblies of all core species. GRs were not annotated for *Aedes albopictus*, for which a partial GR list exists elsewhere^108^. All genomes were annotated using a homology-based iterative BLAST approach^109^. To identify chemosensory receptor genes annotated with the automated GNOMON pipeline, tblastn was used with an e-value cutoff of 1 x 10^−6^ based on gene family-specific query libraries comprised of published mosquito OR, IR, and GR sequences^56^. GNOMON annotations identified this way were individually curated in Geneious Prime (v2025.1.1) and corrected if required. To verify predicted exon-intron boundaries, exonerate (v2.4.0) was used with percent function set to 25^110^. The mosquito-specific query libraries comprised of previously annotated genes were used for the initial round. All validated and corrected GNOMON annotations were then added to the curated query sets and iteratively used to annotate all genomes for chemosensory receptor genes missed by GNOMON using homology-based prediction of exon-intron boundaries in exonerate with percent set to 60 on the assembled end-to-end chromosome scaffolds for each species. Again, all annotations were inspected manually and corrected if needed. The annotation pipeline including all scripts used to parse outputs are available on GitHub (https://github.com/pbrec/csgAnnotationPipeline/tree/main).

### Phylogenetic Reconstruction of Chemosensory Receptor Gene Families

For each chemosensory receptor gene family, a maximum likelihood (ML) gene tree was estimated using IQ-TREE 2^111^ (v2.4.0) to infer their genealogical histories. To this end, gene family-specific protein sequence alignments were constructed using MAFFT^112^ (v7.526) applying the L-INS-i algorithm (--localpair) with –-maxiterate set to 1,000 and –-reorder. ML trees were inferred from the amino-acid alignments with IQ-TREE 2 and automatic substitution-model selection by ModelFinder (-m MFP) and 1,000 ultrafast bootstrap replicates (-B 1000), with the thread count set automatically (-T AUTO). Gene trees were built in this way for all three families (ORs, IRs and GRs). Trees were visualized using TreeViewer^113^. Each chemoreceptor family tree was then manually inspected to identify chemoreceptor branches that contained representative genes from all blood-feeding species, and none from the non-blood-feeding ones.

### Analysis of sexually dimorphic head gene expression

RNA sequencing output (fastq files) was first assessed using FastQC^104^ and inspected for the presence of adapters. Where adapter read-through was present, libraries were trimmed with Trimmomatic v0.39^114^ in paired-end mode using ILLUMINACLIP with the NexteraPE-PE adapter file (seed mismatches 2, palindrome clip threshold 30, simple clip threshold 10, minimum adapter length 2, keepBothReads TRUE), followed by LEADING:3, TRAILING:3, MINLEN:36 and HEADCROP:1. Resulting files were then reanalyzed with FastQC to confirm complete removal of adapters.

Reads were then aligned to the RefSeq genome assembly of the corresponding species using HISAT2^115^ v2.2.1 with 32 threads (-p 32) and otherwise default parameters: hisat2 –p 32 –x “species_index” –1 <sample_R1.fastq> –2 <sample_R2.fastq> –S <sample.sam>. SAM output was converted to coordinate-sorted, indexed BAM using SAMtools^116^. BAM files were counted using HTSeq-count^117^ against the RefSeq annotation of the corresponding species, using the following parameters: –-format bam –-order=pos –-stranded=reverse –-mode=intersection-nonempty –-idattr=gene.

Raw counts were then analyzed separately for each species using DESeq2^107^. For each species, a sample table listing the four female and four male head count files was used to build a DESeqDataSet with DESeqDataSetFromHTSeqCount and the design ∼ condition, where condition was a two-level factor with the male head samples set as the reference level, so that positive log2 fold changes correspond to female-biased expression. Differential expression was tested with DESeq() using the default Wald test, DESeq2’s default independent filtering and Benjamini–Hochberg correction; results were extracted with results() and ordered by adjusted p-value. Genes were called sex-biased only if they satisfied both an adjusted p-value below 0.05 and a linear fold change of at least 1.5 (see below).

### Domain and functional categorization of sexually-dimorphic expressed genes

*Functional categorization of sex-dimorphic head expression*. Per-species differential-expression results for head tissue were restricted to protein-coding genes using the gene biotype field of the corresponding NCBI RefSeq feature table and annotated with a representative RefSeq product description. Protein-domain annotation was taken from InterProScan output filtered to Pfam signatures. Each gene was then assigned to one of 98 functional categories, organized into 18 broad groups, using a curated rule set defined primarily by Pfam accession (350 accessions in total) with regular-expression fallbacks against the RefSeq description for 14 categories that Pfam covers poorly (e.g., mucins, neuropeptides, vitellogenin, doublesex). Where a gene matched several rules, the highest-priority rule was chosen, with ties broken by rule order (see Extended Data^36^).

*Clade contrast and figure construction*. To compare blood-feeding (BF, n = 4) and non-blood-feeding (NBF, n = 3) species, the sex-biased rate of each category was computed per species as the number of biased genes divided by the number of protein-coding genes tested in that category, averaged across species within each clade. *Wyeomyia smithii,* the variable blood-feeding species, was excluded from the contrast. Effect sizes were expressed as Cohen’s h, h = 2(arcsin√p_BF − arcsin√p_NBF), computed separately for the female– and male-biased directions from the same significance criteria used throughout. Categories were eligible for ranking only if they fell outside the “Housekeeping / regulatory” and the unassigned “No Pfam”/“Other Pfam” bins, and if they contained at least three biased genes pooled across BF and NBF species with hits in at least two species.

### Synchrotron scans and rendering

*Volumetric data acquisition.* Following ice anesthesia, samples were transferred to 0.5 mL tubes containing absolute ethanol. The narrow tube diameter restricted sample rotation during acquisition, minimizing sub-micrometer movement. Additionally, using absolute ethanol instead of 70% ethanol prevented the formation of air bubbles during X-ray exposure. Phase-contrast tomographic scans were performed at the ID19 beamline of the European Synchrotron Radiation Facility (ESRF, Grenoble, France. Project number LS-3255. 19.6 keV, 2,600 radiographic projections over an angular range of 180°, pixel size 0.65 µm) or at the SYRMEP beamline of the ELETTRA synchrotron (Project number 20230106; 16 keV, 1,800 radiographic projections over an angular range of 180°, pixel size 0.9 µm). Tomographic slices were reconstructed by conventional filtered back-projection using the SYRMEP Tomo Project (STP) software developed by the SYRMEP team, or using in-house procedures at the ESRF. Images were pre-processed in ImageJ, and reconstructed volumes were segmented in Dragonfly software (v2025.1, build 2063) which was also used to measure structure volumes in voxels. Voxel volumes were normalized to account for the differing pixel size of the two beamlines.

*Structures analyzed.* Volumes were obtained for 11 mosquito species (72 individuals: 42 females, 30 males). For the analyses reported here, nine brain regions (antennal lobe, optic lobe, mushroom body, calyx, protocerebral bridge, fan-shaped body, ellipsoid body, noduli, and mechanosensory/motor neuropil) and three muscles (cibarial pump, head muscle, and clypeal muscle) were measured. Distinct anatomical landmarks make these brain regions straightforward to identify: the antennal lobes are defined by their characteristic glomeruli, while the optic lobes – comprising the lamina, medulla, lobula, and lobula plate – are distinguished by their lateral position and clear regional boundaries. Along the midline of the central protocerebrum, the ellipsoid body and fan-shaped body are readily recognizable by their signature geometries: a bean-like shape for the ellipsoid body and an open-fan contour for the fan-shaped body. Ventral to the ellipsoid body lie the noduli, appearing as two small, dense, elongated structures. Posterior to the fan-shaped body, the protocerebral bridge is identified as a thin, dense tract spanning the two hemispheres. The paired, Y-shaped mushroom bodies stand out as dense neuropils clearly distinct from the surrounding brain tissue; in certain species, their three individual lobes can even be resolved. In the dorso-posterior brain, the mushroom bodies terminate in a discrete region known as the calyx, which exhibits a texture distinct from both the rest of the mushroom body and adjacent neuropils. Following the segmentation of these regions, the mechanosensory neuropil is identified ventral-posterior to the antennal lobes as the region receiving the antennal nerve. Finally, the lateral horn is assigned as the discrete dorso-lateral region situated between the antennal lobe and the mushroom body. The mechanosensory neuropil and lateral horns were measured but excluded from all models and figures because they lacked clear morphological borders. The mushroom body and the calyx measurements were combined to represent the whole mushroom body structure. Total brain volume entered the analyses both directly and indirectly: it served as the size covariate for muscles, as the basis of the size covariate for brain regions (rest-of-brain, defined as total brain minus the tested region), and as the denominator of the ratios plotted in the figure.

### Phylogenetic model and analysis

For each structure and sex, the effect of blood-feeding was tested by phylogenetic generalized least squares (PGLS) on species means. Individual volumes and their corresponding size covariate were log10-transformed, restricted to positive values, and averaged within species, so that each species contributed one observation per sex (a geometric mean) and within-species measurement error was not propagated. The model log_10_(structure) ∼ log_10_(size) + blood-feeding status was fitted with the residual covariance fixed to the Brownian-motion expectation obtained from a time-calibrated maximum-clade-credibility phylogeny pruned to the 11 tested species, where the covariance between two species equals their shared root-to-most-recent-common-ancestor path length (Pagel’s λ fixed at 1). Coefficients were obtained by generalized least squares, and the blood-feeding coefficient was tested with a two-sided t-test on n – 3 degrees of freedom, yielding the reported p-value. p-values were corrected within brain regions and within muscles, females only (males uncorrected) using the Benjamini-Hochberg procedure to yield q-values. *Wyeomyia smithii*, the variable blood feeder, was excluded from every fit so that no reported statistic depended on its placement. The time-calibrated maximum-clade-credibility phylogeny used here was taken from Soghigian et al.^19^. Tomographic reconstruction used the SYRMEP Tomo Project (STP) software or in-house ESRF procedures. Image pre-processing used ImageJ. Statistical analyses and figures were produced in Python using NumPy, pandas, SciPy (t-tests), DendroPy^118^ (phylogeny handling), and Matplotlib.

### Mosquito illustrations

Illustrations of mosquitoes were created by the artist Tatiana Gandlin (https://tatigandlin.com/), based on photographs of live and frozen mosquitoes taken in-house, together with publicly available photographs of each species.

### Statistics, code, and data visualization

Unless stated otherwise above, analyses were run in Python (≥3.8) with pandas, NumPy and SciPy, and in R. Plots were generated with Matplotlib and seaborn (Python) or ggplot2 and patchwork (R); single-nucleus visualizations used Scanpy. Minimum package versions are listed in the requirements and environment files deposited with the code (Extended Data^36^, 00_code). Figures were assembled and finalized in Adobe Illustrator 2023 and 2026 (v27.0 and v30.3).

### Data availability

The seven genome assemblies generated in this study and accompanying RNA sequencing have been deposited in NCBI databases and are released openly for reuse. The assembly accession and annotation release for each species are listed below. Assembly data and RNA sequencing for all species except *Sabethes cyaneus* are available on BioProject PRJNA942918. *Sabethes cyaneus*’ data are available on PRJEB53261. *Aedes aegypti*’s midgut RNA-seq dataset is available on PRJNA1502948. Publicly available datasets reanalyzed here are *Aedes aegypti*’s single-nucleus RNA-seq^49^, bulk RNA-seq of anterior and posterior midgut^48^, and ovaries across the blood-feeding and egg-laying cycle^55^.

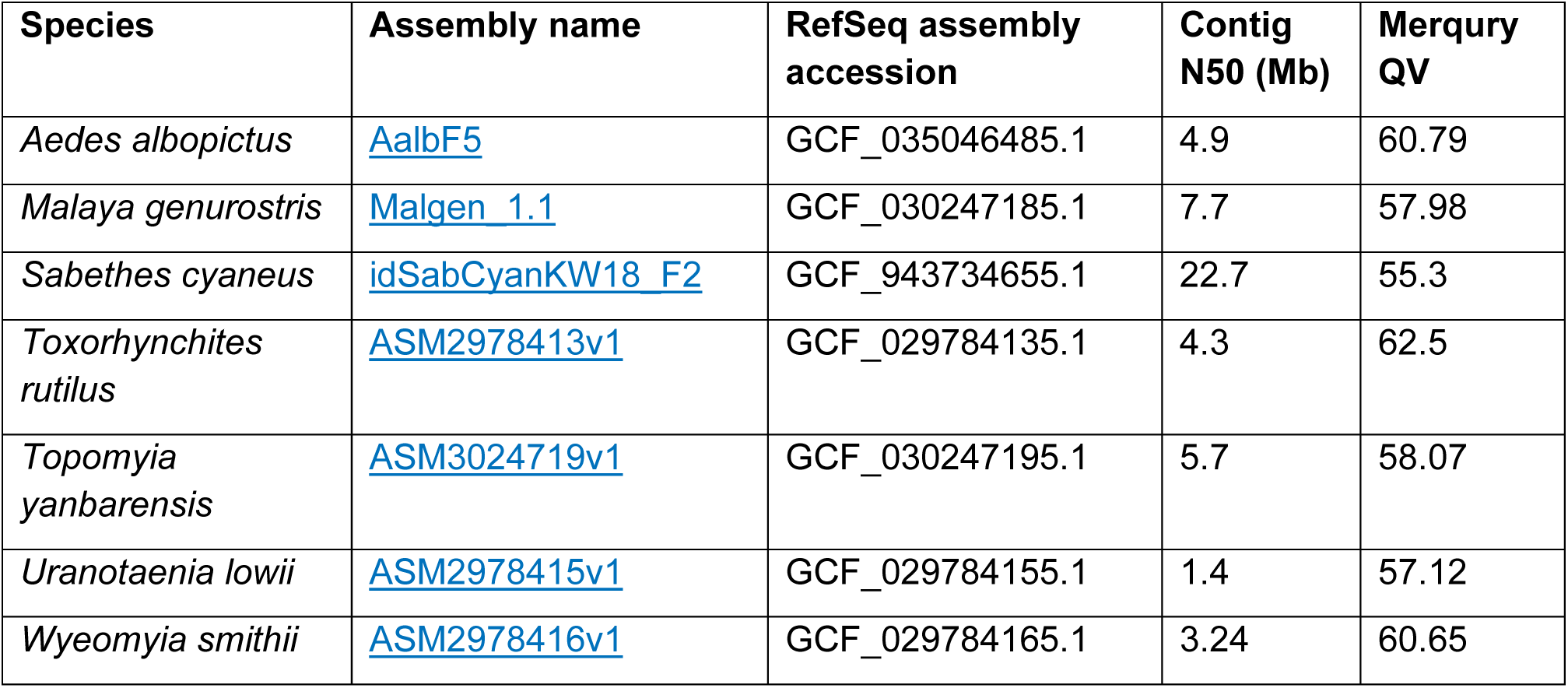

All processed data, annotations and analysis code generated in this study are available in the Extended Data on Zenodo at https://doi.org/10.5281/zenodo.22646228, organized as follows: RepeatMasker summaries (01_repeats); endogenous viral element statistics and the viral protein database (02_EVEs); InterProScan output and the TE-like genes removed before orthology analysis (03_interproscan); OrthoFinder hierarchical orthogroups and species tree, the 25 blood-feeding-specific gene clades, and the gene-loss validation tables, microsynteny anchors and per-locus plots (04_orthology); manual OR, IR and GR annotations (GFF3), gene trees and per-species counts (05_chemoreceptors); DESeq2-normalized count matrices and gene rankings for the midgut time course and the reanalyzed midgut and ovary datasets (06_bulk_rnaseq); per-species head female-versus-male DESeq2 results and the functional-category rules (07_head_sexbias); antennal cluster sex composition derived from the Mosquito Cell Atlas (08_snRNAseq); and brain-region and head-muscle volumes for all 72 individuals, the time-calibrated phylogeny and PGLS results (09_tomography). Every file is listed with an md5 checksum in MANIFEST.tsv. Raw synchrotron microtomography data are available from the ESRF data portal (https://data.esrf.fr/investigation/1305995198/datasets) and in Extended Data^36^.

### Code availability

All custom code is archived in Extended Data^36^ (00_code). This includes the EVE-classifier detection and analysis pipeline; the pipeline for functional categorization of sexually dimorphic head gene expression; the Scanpy scripts for single-nucleus RNA-seq visualization; and the phylogenetic generalized least squares analysis of brain and muscle volumes. Claude (Anthropic) and Gemini (Google) were used to assist with code, plotting and data-consistency checks, all of which were reviewed and verified by the authors.

